# Holobiomes in succession: post-glacial microbial communities are structured by hosts, time and habitat heterogeneity

**DOI:** 10.64898/2026.01.20.700596

**Authors:** Adam Taylor Ruka, Vojtěch Lanta, Samresh Rai, Kateřina Čapková, Thinles Chondol, Inga Hiiesalu, John Davison, Lucie Vančurová, Jan Kučera, Jiří Doležal, Roey Angel, Klára Řeháková

## Abstract

1. Glacier forefields in the high-desert region of Ladakh (northwestern Himalaya) are colonized by a variety of interdependent organisms, including lichens, prokaryotes, fungi, mosses, and vascular plants, along a successional gradient. Together with bulk soil, these hosts and their associated microorganisms form a broader microbial metacommunity (holobiome) whose structure, interactions and functions remain poorly underexplored in one of the Earth’s most extreme and climate-sensitive environments.
2. Using a multidisciplinary approach combining glacial chronosequence transects, GIS-derived topographic variables, soil properties, and plot cover measurements, we assessed the abiotic and biotic factors influencing bacterial and fungal communities sequenced from different hosts and bulk soil (hereafter sources). Microbial composition was primarily shaped by source identity, though certain sources, such as biological soil crusts (BSCs), mosses, and plant rhizospheres, also showed relationships with moraine age in either bacterial or fungal communities.
3. Bacterial and fungal community congruence was tested using Procrustes analyses, revealing that mosses maintained tightly coupled inter-kingdom relationships throughout the glacier forefields. However, the degree of congruence in plant rhizospheres and bulk soils was influenced by topographic variation and moraine age, respectively.
4. Co-occurrence network analyses revealed that early successional microbial communities were assembled more stochastically, with bacteria being more interconnected than fungi. In contrast, late successional stages were more compartmentalized, being more structurally stabile, likely driven by increased plant cover and functional redundancy among microbial taxa.
5. Keystone bacterial and fungal taxa were identified in plant rhizospheres and bulk soil using a dual-criteria approach related to inter-kingdom congruence and network node eigenvalues. Furthermore, some of these taxa were associated with environmental factors, suggesting topographic heterogeneity and successional age can promote or deter the influence of keystone taxa.
6. *Synthesis:* This study reveals the impact of both macroorganism colonization (i.e. plants, mosses, and lichens) and microcommunity establishment (BSCs and bulk soil), as abiotic and biotic sources, on microbial metacommunity assembly in glacier forefields. By adopting a broader approach across different spatial scales, we demonstrate that while plant colonization plays a central role in shaping microbial metacommunities, its effects are modulated by topographic variation along the chronosequence.

## 1. Introduction

An estimated 142,000 km^2^ of habitat around the margins of glaciers in alpine and polar regions will be newly exposed by the end of this century due to accelerated warming (Graversen et al. 2008, Pepin et al. 2015, Zimmer et al. 2022, Nigrelli and Chiarle 2023). These extreme environments, characterized by a rocky, nutrient-poor soil and cold, windy conditions, are subsequently colonized by tolerant, pioneer organisms, including microorganisms, cryptogams (mosses and lichens), and vascular plants (Bueno de Mesquita et al. 2017, Darcy et al. 2018, del Moral et al. 2021). To survive in such environments, colonizing organisms must employ different persistence strategies, such as chemolithotophy, atmospheric nitrogen fixation, freezing and desiccation tolerance, and inter-kingdom symbioses. These adaptive mechanisms enable a diverse array of species to establish themselves in the harsh conditions of glacier forefields. Thus, a mosaic of macroorganisms and barren soil establishes within a glacier forefield and provides unique microniches for their associated microbiomes (Bueno de Mesquita et al. 2017, Touchette et al. 2023, Wicaksono et al. 2024). For example, vascular plants host specific root microbial communities through selective processes, such as root exudates in the rhizosphere (Ciccazzo et al. 2014). Cryptogams (lichens and mosses) and biological soil crusts (BSCs) further host specific microbial consortia through increased nutrient availability (Cardinale et al. 2006) and interdependent ecological relationships (Cania et al. 2020). Meanwhile, the barren soil substratum in glacier forefields primarily serves as a reservoir of microbial diversity, from which the hosts can recruit their respective microbial communities (Chen et al. 2023). In conjunction with atmospheric and erosional deposition as sources of microbial colonization (Elster et al. 2007, Franzetti et al. 2020, Jensen et al. 2022), these various biotic and abiotic compartments (collectively referred to as ’sources’ hereafter) constitute a microbial metacommunity (or holobiome), shaped by positive and negative interactions between bacteria and fungi at small spatial scales (Deveau et al. 2018). Yet, despite numerous studies assessing microbial assembly processes in glacier forefields (Hodkinson et al. 2003, Vimercati et al. 2022, Mandolini et al. 2025), disentangling the relative importance of these sources, such as plants, cryptogams, BSCs, and bulk soil, across environmental gradients remains unexplored.

Metacommunity theory posits that microbial communities assemble through a combination of deterministic and stochastic processes, with their relative influence varying across ecological contexts and geographic scales (Leibold et al. 2004, Vellend 2010, Miller et al. 2018). Deterministic processes, consistent with niche theory, describe community assembly as being governed by environmental selection; being driven by abiotic (e.g., pH, soil nutrients, temperature) and biotic factors (e.g., host identity, biological interactions) which typically shape patterns at larger spatial scales (Riddley et al. 2025). Stochastic processes follow neutral theory, attributing community structure to random events such as phylogenetic drift and dispersal limitation, and dominate at smaller geographic scales, where habitat heterogeneity and environmental filtering exert less influence (Germain et al. 2021, Yang et al. 2024). Additionally, there is some evidence that the importance of these processes can differ among microbial groups, with deterministic processes more strongly influencing bacterial communities (Riddley et al. 2025), while stochastic processes appear to impact fungal community structure (Mandolini et al. 2025). However, within a glacier forefield, habitat heterogeneity is pronounced, with predictable changes in soil development and vegetation cover (Matthews 1992, Bradley et al. 2014). These successional patterns suggest microbial community composition should be heavily impacted by specific changes in abiotic and biotic microniches, such as the occurrence of macroorganisms and bulk soil fertility, which act as hosts or reservoirs of organisms.

Across a glacier chronosequence, the colonization and dominance of higher organisms rely upon substrate availability and microclimatic conditions. Upon newly-exposed rock surfaces, lichens are among the first to colonize following glacier retreat, contributing to primary production and nutrient cycling through their symbiotic phototrophs and associated microbiota (Cornelissen et al. 2007). Apart from their obligate phototrophic partners, lichen thalli host diverse microbial taxa that form biofilm matrices while enhancing nutrient availability and conferring resistance to abiotic stress (Grube et al. 2009). Concurrently, BSCs establish on fine mineral substrates in recently deglaciated zones, composed of tightly interdependent communities of phototrophic and heterotrophic microorganisms. These organisms form a biofilm matrix and perform key metabolic functions such as photosynthesis and respiration, contribute to soil carbon and nitrogen pools, promote substrate stabilization through the production of extracellular polysaccharides (Belnap et al. 2003, Chamizo et al. 2012), while also associating with fungal counterparts (Liu et al. 2024). Mosses are also able to colonize these glacial habitats, particularly in moist microclimatic niches, at an extremely early stage, sometimes even growing upon glacier ice (Dickson and Johnson 2014, Anderson et al. 2025). Given the slower decomposition rate and distinct tissue composition (Lang et al. 2009), mosses influence microbial community composition and may serve as an understudied reservoir for microbial diversity (Bueno de Mesquita et al. 2017). In contrast, vascular plants increase in prevalence over successional time and profoundly alter microbial communities through root exudation, shading, littering, and active recruitment of mutualistic microbial taxa (Ciccazzo et al. 2014, Kuzyakov and Blagodatskaya 2015, Kawasaki et al. 2021). Concomitantly, this increase in plant coverage coincides with increased soil carbon and nutrient availability, leading to changes within bulk soil microbial communities (Philippot et al. 2011, Ruka et al. 2023). Nevertheless, across environmental gradients, these colonizing organisms encounter varying levels of stress, which in turn modulate bacterial–fungal interactions and influence overall community assembly. Although inter-kingdom relationships between bacteria and fungi are not well understood, evidence shows bacterial endosymbionts and physical attachment to fungi (Steffan et al. 2020). This suggests that keystone microbial taxa associated with specific sources are likely to influence co-occurring microbes from other kingdoms, leading to greater congruence or coupling between communities.

The chronosequence approach operates on the key assumption that sites within a glacier forefield undergo succession with comparable initial environmental conditions, thereby allowing researchers to infer that older sites represent the future states of younger ones (Wojcik et al. 2021). Ecological succession theory suggests that these autogenic factors (e.g. biota, interactions, soil development) work in tandem with allogenic factors (e.g. erosion, extreme weather events, landslides) to establish the successional trajectory of a given environment (Matthews 1992). Specifically within glacier forefields, considerable heterogeneity in topographic features, such as slope, aspect, and concavity, can significantly influence microclimatic conditions and resource availability, consequently shaping the colonization dynamics of microbes, cryptograms and vascular plants (Solon et al. 2021, Ruka et al. 2023). These microenvironmental variations interact with broader climatic changes, as shifting climate regimes in alpine and polar regions have altered the extent and duration of seasonal snow cover, with some alpine areas experiencing an increase of up to 43 snow-free days over the past two decades (Notarnicola 2020). Consequently, these changes add to the potential growing season and increase recruitment of vascular plants populations in snow-free locations (Jandova et al., 2025). Still, another consequence of climate change is the higher frequency of extreme precipitation events, including snowfall at high elevations during the growing season, which can negatively impact plant performance (Jandova et al., 2025) and potentially decouple plant–microbial interactions. Despite growing recognition of these changes, a comprehensive assessment of how autogenic factors (time since deglaciation, sources) and allogenic drivers (topography, climate) shape microbial community assembly across glacier forefields remains lacking.

The heterogeneity of glacier forefields, shaped by both abiotic and biotic factors, plays an important role in microbial community assembly across small and large geographic scales. Although numerous studies have explored broad changes in soil microbial communities using chronosequence approaches (Garrido-Benavent et al. 2020, Vimercati et al. 2022, Ruka et al. 2023, Acuña-Rodríguez et al. 2023), the distinct contribution of different sources, especially in conjunction with environmental heterogeneity, has not been investigated. Therefore, in this study, we aimed to characterize bacterial and fungal communities associated with distinct hosts and the bulk soil (sources) along a glacial successional gradient in the northwestern Himalaya region of Ladakh, India. Considered to be a “third pole”, comparable to the Arctic and Antarctic in freshwater availability and glacial extent (Banerjee et al. 2021), the Himalaya represent an understudied region for microbial biodiversity. Therefore, this study follows previous studies in the region revealing the upper elevational distribution limit of plants (Dolezal et al. 2016), diversity patterns of plant-symbiotic arbuscular mycorrhizal fungi at high elevations (Hiiesalu et al. 2023), and differing microbial successional pathways based on elevation and climate (Ruka et al. 2023). To further the understanding of successional processes and microbial metacommunity assembly in the region, we employed a multidisciplinary approach that integrated in-situ cover measurements, molecular sequencing of bacteria and fungi, soil physicochemical analyses, and geographic information system (GIS) tools to quantify topographic variables and infer potential shifts in climatic regimes. Through network analyses and Procrustes tests, we examined inter- and intra-kingdom relationships to identify the dominant factors influencing microbial metacommunity assembly and keystone taxa in the highest elevation glacier forefield chronosequences known to be studied. Thus, we hypothesized the following: 1) bacterial and fungal community composition will differ depending on the source (lichen, moss, BSC, plant rhizosphere, and bulk soil; 2) within each source, variation in topographic features is expected to influence inter-kingdom congruence between bacterial and fungal communities, with moraine age playing a lesser role than other topographic factors for microclimate-sensitive organisms such as mosses and vascular plants; 3) microbial metacommunity assembly will be strongly influenced by the presence or absence of plants, due to their selective effects on microbial communities and their role in promoting soil development compared to other sources.

## 2. Methods

### 2.1. Study sites

The study was performed in the Ladakh region of India, a high-altitude desert located in the northwestern ranges of the Himalaya. Overall, the region often receives less than 100 mm year^-1^ of total annual precipitation due to a strong rain-shadow effect related to orographic barriers. With the region being the intersection of multiple mountain ranges (e.g. Karakoram, Pangong, Greater Himalaya ranges), bedrock material varies considerably in the region from granitic and gneissic, while limestone is common at higher elevations (Phillips 2008, Kowser et al. 2017). Mean annual air temperatures in the regions vary from 1.6 to 10.4 °C while mean annual soil temperature ranges from 0.3 to 7.4 °C (Ruka et al. 2023). The two study locations, Tso Moriri (32°59′N, 78°24′E) and Nubra (34°45′N, 77°35′E) valleys, are separated by 200 kilometers, yet both are near the transition to the Qinghai-Tibetan Plateau. Additionally, they differ in elevation, with glacier tongues terminating in Nubra at 5,100-5,400 m.a.s.l. while in Tso Moriri they terminate at 5,800 m.a.s.l. Previous studies at the sites investigated environmental stressors and soil phototrophs (Rehakova et al. 2019), plant-facilitation (Dvorský et al. 2013), and plant recruitment (Jandova et al., 2025). Notably, glacier forefield successional pathways of plants and soil bacteria communities were studied at the two locations, finding similar bacterial taxa inhabited the environments closest to the glacier tongue (Ruka et al. 2023).

### 2.2. Field work

Transects were performed at four glacier forefields in the Ladakh region during the 2022 field season (15/8/22-29/8/22). Three glacier tongues of the South Shupka Kunchhang Glacier in Nubra (5400, 5300, and 5150 m) were selected for the study due to geographic proximity, while one glacier tongue of Chamser Glacier in Tso Moriri (5800 m) was selected. Starting at the terminus of each glacier tongue, nine plots were established across three moraines with three plots per moraine (n = 36). Plots were established along the transect at the front, top and back side of the moraine to account for topographic variation. Relative age of the moraines was categorized as young, intermediate, and old based on sequential moraines away from the glacier tongue (Figure 1). Within each 1 m^2^ plot, cover (%) was recorded for vascular plants, lichens, mosses, soil crust, stones, and bare soil. Furthermore, latitude and longitude coordinates, elevation (m.a.s.l.), and distance from the glacier tongue were recorded. Sampling from each plot included a composite bulk soil sample from three subsamples (0-10 cm), lichens, soil crusts, mosses, and fine root samples of the three most dominant plants within the plot. Due to some plants having minimal fine root material, samples were pooled from at least two individuals. From each plot, 3-10 samples in total were collected depending on occurrence along the transect. Plant species were identified in the field using a local flora guide (Dvorskỳ et al. 2018). All samples were preserved by drying in separate plastic bags with silica gel (Bainard et al. 2010), while bulk soil samples were air-dried after removing a subsample for molecular analyses. All samples were transported to České Budějovice, Czech Republic for further molecular analyses, identification of mosses and lichens, and soil physicochemistry analyses. Moss and lichen identification methods are detailed in the supplementary material (Appendix 1).

**Figure 1.**
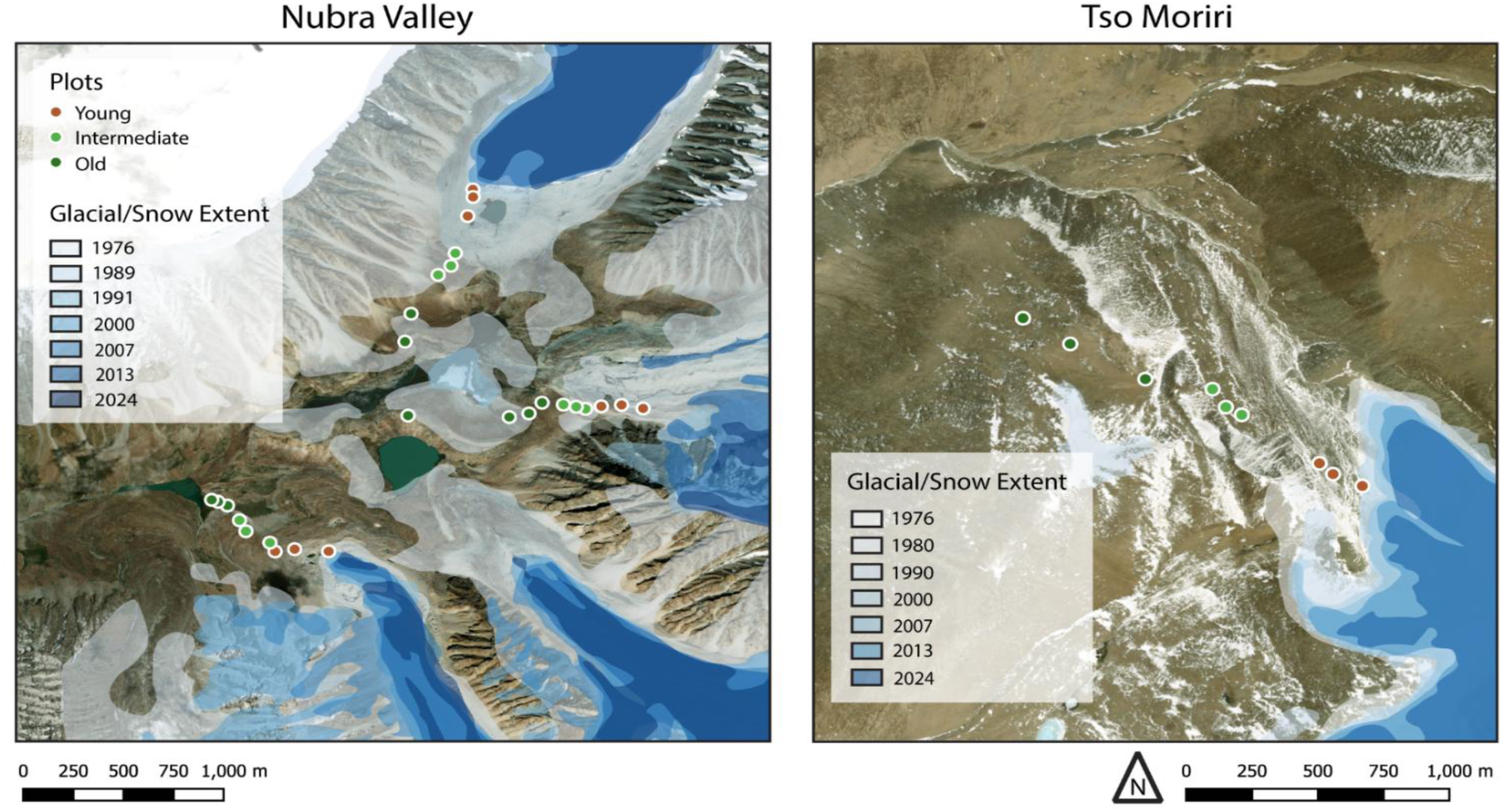
Maps of the studied glacier forefields with past glacial/snow extent based on historical satellite imagery retrieved from the USGS Earth Explorer. Images were selected during the period of minimal snow cover in the Ladakh region (August–October) and optimal visibility. Maps were built in QGIS and projected in Pseudo Mercator using WGS 84 coordinate system.

### 2.3. Soil physiochemistry analyses

Air-dried bulk soil samples from each plot were stored at –20 °C until further processing at the Institute of Botany in Třeboň, Czech Republic. Then, soil samples were oven-dried at 80 °C and 2 mm sieved before being ground in a mortar. Weight measurements before and after sieving were used to calculate texture (% > 2 mm). Soil pH was measured in a 1:5 (soil:deionized water) suspension. Soil organic matter content (OM, %) was determined by dry combustion at 450 °C for 5 hours. Total nitrogen (TN; mg kg^-1^) was quantified using a Lachat QC 8500 automated injection analyzer (three-channel FIA system), with signal detection via the Thermo Scientific Eager Xperience platform (Eagersmart 2016) (Sparks 1996). Total phosphorus (TP; mg kg^-1^) was analyzed according to U.S. EPA Method 365.4 and Standard Methods 4500–P G. Major cations (Ca²⁺, Mg²⁺, K⁺, Na⁺) were measured using flame atomic absorption spectrometry (ICP-MS, ICP-QQQ; Agilent Technologies, Tokyo, Japan) (Kopáček et al. 2004).

### 2.5. DNA extraction, PCR amplification, and sequencing of sources and soil

DNA from root, lichen, moss, BSC, and bulk soil samples (240 total) were extracted using DNeasy PowerSoil Pro Kit (Qiagen) after adding 100 μl of phosphate buffer for rehydration. On average, 0.2 grams of material was used for extraction, and DNA concentration was measured using Quant-iT™ PicoGreen™ dsDNA Assay Kit (Thermo). DNA extracts were sent to AsperBiogene in Tartu, Estonia, where PCR amplification, library construction, and high-throughput sequencing were performed on an Illumina MiSeq sequencer (Illumina). For fungal barcoding, the ITS2 region was targeted using the fITS7 (5’-GTGARTCATCGAATCTTTG-3’) (Ihrmark et al. 2012) and the ITS4ngsUni (5’-CCTSCSCTTANTDATATGC-3’) (Tedersoo and Lindahl 2016). For bacteria, the V4 region of the 16S rRNA gene was amplified using the primers 515F-Y (5′-GTGYCAGCMGCCGCGGTAA-3′) (Parada et al. 2016) and 806R (5’-GGACTACNVGGGTWTCTAAT-3’) (Apprill et al. 2015).

### 2.6. Bioinformatics

Raw short-read sequencing data (16S rRNA and ITS2) were processed using a pipeline based on DADA2 (Callahan et al. 2016). Primers and linker regions were trimmed using Cutadapt (V3.5) (Martin 2011), followed by quality filtering, merging, denoising, and chimera removal within DADA2 in R v4.4.1 (R Core team, 2025). Non-standard filtering parameters included: filterandTrim function with maxEE = c(2,2) and pseudo pooling. Taxonomic assignment was performed using the SILVA138.2 database (Quast et al. 2013) for 16S and UNITE v10.0 (Abarenkov et al. 2024) for ITS2 sequences. Heuristic determination was done by applying the decontam R package (Davis et al. 2018), and then compiled to unique sequences based on a 97% identity threshold. Taxonomic filtering removed Archaea, Mitochondria, Chloroplast, and unclassified sequences from the 16S data and Viridiplantae. Alveolata, Heterolobosa, Ichthyosporia, Metazoa, Rhizaria, Stramenopila, and unclassified sequences from the ITS2 data. A minimum library size of 1000 reads, singleton removal, and a prevalence threshold of 1% (three samples) for sequences were applied to both bacterial and fungal datasets. For bacteria, a total of 211 samples remained (mean library size: 20,479) with 13,907 taxa and 4,321,109 total reads. For fungi, a total of 207 samples remained (mean library size: 9,739) with 2,594 taxa and 2,016,152 total reads. Subsequently, phyloseq objects were created (McMurdie and Holmes 2013) for both bacterial and fungal reads using processed sequence tables, taxonomy, and metadata for bacteria and fungi. To account for variability in the ITS2 region, duplicated taxa, and statistical cohesiveness, both phyloseq objects (for the 16S rRNA and the ITS) were agglomerated at the genus level. Alpha diversity metrics (Chao richness and Shannon-Weiner H’ diversity) were calculated using the plot_richness function in the phyloseq package. Last, bacterial and fungal count data were converted to relative abundance values for downstream analyses.

### 2.7. GIS Applications

QGIS 3.42.3 was used for all GIS-derived variables. Topographic variables were extracted using latitude and longitude point features overlaid on the Global Multi-Resolution Topography (GMRT) Synthesis layer, which has a resolution of 30 × 30 m. Topographic Position Index (TPI), slope, and aspect were calculated using the respective tools in QGIS with default settings. TPI indicates the relative position of a location in the landscape, capturing whether it lies in a concave (e.g., valley) or convex (e.g., ridge) terrain. Due to aspect being a circular variable (0° = 360°), it was converted into sine and cosine components to allow for linear statistical analysis. In this transformation, the sine of aspect represents the east–west orientation (positive values = east-facing slopes), while the cosine of aspect represents the north–south orientation (positive values = north-facing slopes).

To assess historical changes in glacial extent and minimum annual snow cover, a geographic database was developed in QGIS by acquiring LANDSAT raster data from the USGS EarthExplorer (U.S. Geological Survey, 2024). GeoTIFF layers spanning from the 1970s to the present were downloaded, selecting one image per decade based on cloud-free conditions and minimal snow cover, typically in September or October. Early imagery from 1976 and 1980 had a resolution of 60 × 60 m, while later images had a resolution of 30 × 30 m. Glacial features were digitized from reflectance bands using the polygon drawing tool in QGIS, followed by two iterations of polygon smoothing. Among these layers, the September 1976 image exhibited the greatest glacial and snow cover extent and was used to generate a categorical variable labeled “Snow 1976.” All spatial layers were projected using the Pseudo Mercator coordinate system (WGS 84 datum).

### 2.8. Statistical Analyses

All statistical analyses were performed in R v4.4.1. As a preliminary step, redundancy analyses (RDA; R package: vegan) were carried out to evaluate the effect of environmental variables (moraine age, TPI, slope, aspect [sine and cosine], Snow 1976, and elevation) on two sets of response variables: (1) root-mean-squared soil physicochemistry parameters, and (2) isometric log-ratio transformed cover (%) measurements, separately. Each RDA was conducted separately for these two response sets, with models conditioned by location to account for spatial variation and identify broader regional trends. The statistical significance of the full models and individual predictor terms was tested using permutation-based ANOVA with 999 permutations.

Bacterial and fungal alpha diversity metrics (Shannon-Weiner diversity index, richness) were analyzed using linear mixed-effect models (R package: lmer4) with moraine age (young, intermediate, old), source (lichen, soil crust, moss, bulk soil, plant), and location (Tso Moriri and Nubra) as categorical fixed effects. To account for multiple sources being collected from each plot, the plot ID was used as a random effect. Model significance was assessed using analysis of variance via Type II F-tests (R package: car). Post-hoc comparisons of alpha diversity among sources within each moraine age were conducted using aligned rank transformation ANOVA (R package: ARTool) followed by pairwise comparisons of estimated marginal means (R package: emmeans), with false discovery rate (FDR) correction applied to p-values. The aligned rank transformation was applied as a non-parametric approach due to unequal sample sizes within moraine ages. Then, bump plots were created to visualize the unique vs. shared genera from each source across moraine ages. Subsequently, the genera shared by all sources across all moraine ages was termed the ‘core’ community and used for network analyses and Procrustes analyses.

For microbial beta-diversity analyses, a center log-ratio (CLR) transformation was applied with a pseudo count of 1e^-6^ added (apart from network analyses where a pseudocount of 1 was added). A PCoA was performed on the CLR-transformed community data using Euclidean distance for both bacteria and fungi (R package: microViz). Beta-dispersion (distance to centroid) was calculated from the resulting ordinations (R package: vegan), followed by linear mixed-effect model tests as described in the previous paragraph. Permutational multivariate analysis of variance (PERMANOVA) analyses (999 permutations) were paired with PCoA analyses using moraine age, source, and location as explanatory variables and permutations restricted to within plots (strata; R package: vegan). Upon finding that source identity explained the most variation in bacterial and fungal community composition, subsequent PERMANOVA analyses were performed on separate sources with moraine age and location as explanatory variables. Plot ID was included as a fixed effect for sources which had multiple samples per plot (lichens, mosses, and plants). P-values were adjusted using FDR correction for multiple comparisons. Last, composition bar plots were generated based on relative abundance of pooled samples of all shared taxa and individual sources.

Patterns of coupling or decoupling between bacterial and fungal communities across environmental gradients were assessed using Procrustes analysis (R package: vegan) (Peres-Neto and Jackson 2001). To account for potential spatial or location-specific variation in community composition, principal component analyses (PCA) were performed separately for fungal and bacterial communities using CLR-transformed relative abundance data, while conditioning by sampling location. The ordination scores from the conditioned PCAs served as input matrices for Procrustes analysis, which was conducted with 999 permutations and symmetric scaling (symmetric = TRUE). This analysis was applied to the full dataset (all samples combined) and separately to each source. The resulting Procrustes M² value, a measure of the dissimilarity between two ordination configurations, where lower values indicate stronger concordance, was used to quantify the degree of community coupling. For each Procrustes analysis, residual distances were also extracted at the sample level and evaluated in relation to environmental variables using linear mixed-effects models, using plot ID as a random effect where applicable. Distance from the glacier tongue (in meters) was used as a proxy for a continuous moraine age variable. To identify taxa potentially associated with bacterial–fungal community decoupling, Spearman rank correlations were calculated between individual taxon abundances and Procrustes residuals.

To assess microbial co-occurrence patterns across individual plots, genus-level fungal and bacterial network analyses were performed separately (R package: NetCoMi). For each plot, samples were subset using the ‘core’ community described previously, ensuring a minimum of three samples per analysis. Plots with fewer than three samples were excluded from the analysis. To prevent a bias upon global network and node parameters due to a higher number of input samples, subsetting was performed on plots with more than four samples by randomly selecting four samples without replacement in ten independent iterations. Networks were constructed for individual plots using Spearman correlations on CLR-transformed abundance data, with zero replacement via a pseudocount (1). For fungal networks, all taxa were included, while bacterial networks included 200 taxa based on the highest frequency among samples. Sparsification was performed using the default correlation threshold (0.3). The resulting networks were analyzed using a hierarchical clustering method to compute global topological parameters (average node dissimilarity, average path length, clustering coefficient, modularity, edge connectance, natural connectance, network density, perturbation effect parameter) and node-level centrality measures (degree, betweenness, closeness, eigenvector centrality, and clustering coefficient). These metrics were aggregated and averaged across subsetted iterations for global and node-level parameters. Keystone taxa from the network analyses were selected based upon the criteria of an eigenvalue > 0.95. To assess how different sources influence network structure, the network construction and analysis described above were repeated while individually excluding each source. Global network parameters from these exclusion treatments were then analyzed using an aligned rank transformation ANOVA (non-parametric approach), with plot ID included as a random effect. Post-hoc pairwise comparisons between source exclusions were performed using estimated marginal means, and p-values were adjusted for multiple comparisons using the false discovery rate (FDR) method.

A hierarchical redundancy analysis (RDA) approach was employed to assess the influence of environmental gradients on bacterial and fungal network structure. Separate RDAs were first conducted for three groups of explanatory variables: (1) environmental variables (TPI, slope, aspect [as sine and cosine], moraine age, snow extent from 1976, elevation), (2) plot cover variables (cover [%] of stones, bare soil, soil crust, moss, lichen, graminoids, and forbs), and (3) soil physicochemical parameters (pH, texture, OM, TN, TP, Ca, Mg, and Na). All RDAs were constrained by sampling location to account for spatial variation and limit the effect of unequal number of transects between the locations. The first and second axis scores from each of these RDAs were extracted and used as representative summaries of each variable group. These scores were then included as predictors in a second-level RDA, performed separately for bacterial and fungal datasets, where the response matrix consisted of root-mean-square (RMS) values of global network parameters derived from microbial co-occurrence analyses. Statistical significance of the overall model, individual terms, and canonical axes was assessed using permutation-based ANOVA (999 permutations). This two-tiered RDA approach enabled partitioning of variance in microbial network structure attributable to environmental, cover, and soil drivers while controlling for site-specific effects.

To identify keystone bacterial and fungal genera associated with environmental gradients in glacier forefields, we employed a dual-criteria approach. Taxa were first selected based on their consistent identification as keystone species in microbial co-occurrence networks (node eigenvalue > 0.95). In parallel, indicator taxa were identified through significant Spearman rank correlations (p < 0.05) between their relative abundances and Procrustes residuals, with a maximum relative abundance threshold of >1%. Genera meeting both criteria were considered environmentally relevant keystone taxa. These overlapping genera were then ranked according to the number of network analyses in which they were classified as keystone taxa. For the top ten genera, Spearman rank correlation tests were subsequently performed to assess relationships with key environmental variables identified in the Procrustes analyses.

## 3. Results

### 3.1. Environmental variables, cover measurements, and soil physicochemistry variables

Preliminary RDAs showed a predominant effect of moraine age (p <0.01) upon both soil physicochemistry parameters and cover measurements from the studied transects (Supplementary Figure 1, 2). The amount of constrained variation was higher for cover measurements (50.2%) than for soil parameters (34.6%). However, the amount of conditioned variation by location was higher for soil parameters (23.8%) than for cover measurements (5.4%). On old moraines, graminoid and forb cover increased with higher OM, TN, and other micronutrients such as Ca, K, and Mg. Young moraines had higher coverage of stones and coarse-grained soil texture. Among the remaining environmental variables, TPI was often related to the second axis and negatively correlated with soil crust cover, bare soil cover, while being positively correlated with increasing pH. Taken together, these results suggest a stronger difference in soil properties between the two study locations, while cover measurements differed less, although both RDAs demonstrate that moraine age is the primary factor determining their variation.

### 3.2. Abiotic and biotic sources of microbial diversity

Among the five sources, plants were most frequently sampled source along the studied chronosequences (n=94 samples) (Supplementary Table 1). In total, rhizosphere microbial communities were assessed from 31 plant species, including *Carex borii* (11), *Waldhemia tridactylites* (9), *Thylacospermum caespitosum* (8), *Saussarea glacialis* (7), and *Carex sagaensis* and *Potentilla palmirica* (both), as the most frequently sampled species. Lichens were the second most common sampled source (7 fungal hosts identified; 36 total) with *Xanthoria elegans* (6) and *Gyalolechia bracteata* (4) being particularly prevalent. BSCs followed with 36 successfully sequenced samples. Mosses were second to last with 28 microbial communities sequenced and 10 moss species identified (*Distichium capillaceum and Bryum* spp. being the most prevalent). Bulk soil samples (21) were the least successful in sequencing, particularly in early moraine plots, highlighting the low microbial biomass in these soils.

### 3.3. Bacterial and fungal alpha-diversity

Genus-level bacterial and fungal alpha-diversity metrics were primarily influenced by source; however, moraine age also had a significant effect on fungal diversity (Supplementary Table 2). In bacterial communities, both Chao richness and Shannon diversity (H’) were consistently lowest in lichen samples across all moraine ages. In contrast, soil crusts, mosses, bulk soil, and plants exhibited similar diversity in young moraines, while mosses and plants showed higher richness and diversity in intermediate and old moraines (Figure 2A). For fungal communities, Chao richness generally increased with moraine age. Lichens and soil crusts had the lowest fungal Chao richness and H’ diversity across all ages, while plants, mosses, and bulk soil samples showed progressively higher values (Figure 2B). Bump plots illustrating unique and shared genera among sources revealed that shared genera were predominant throughout the glacial chronosequence. The proportion of shared bacterial genera increased from 78% to 84%, and shared fungal genera from 47% to 57%, across moraine ages (Figure 2D). Among sources, plants hosted the highest proportion of unique bacterial genera, while bulk soil, followed by plants, harbored the most unique fungal genera across all moraine ages. Although these patterns suggest substantial exclusivity of microbial genera by source identity, a large fraction of sequencing reads belonged to the ’core’ microbiome, defined as genera shared across all moraine ages. Specifically, the core comprised 95.7% (4,037,426) of bacterial reads and 83.9% (1,677,114) of fungal reads. These findings present both the niche-specific associations of certain taxa and the widespread distribution of others, resulting in distinct patterns of microbial alpha-diversity across glacial successional stages in different sources.

**Figure 2.**
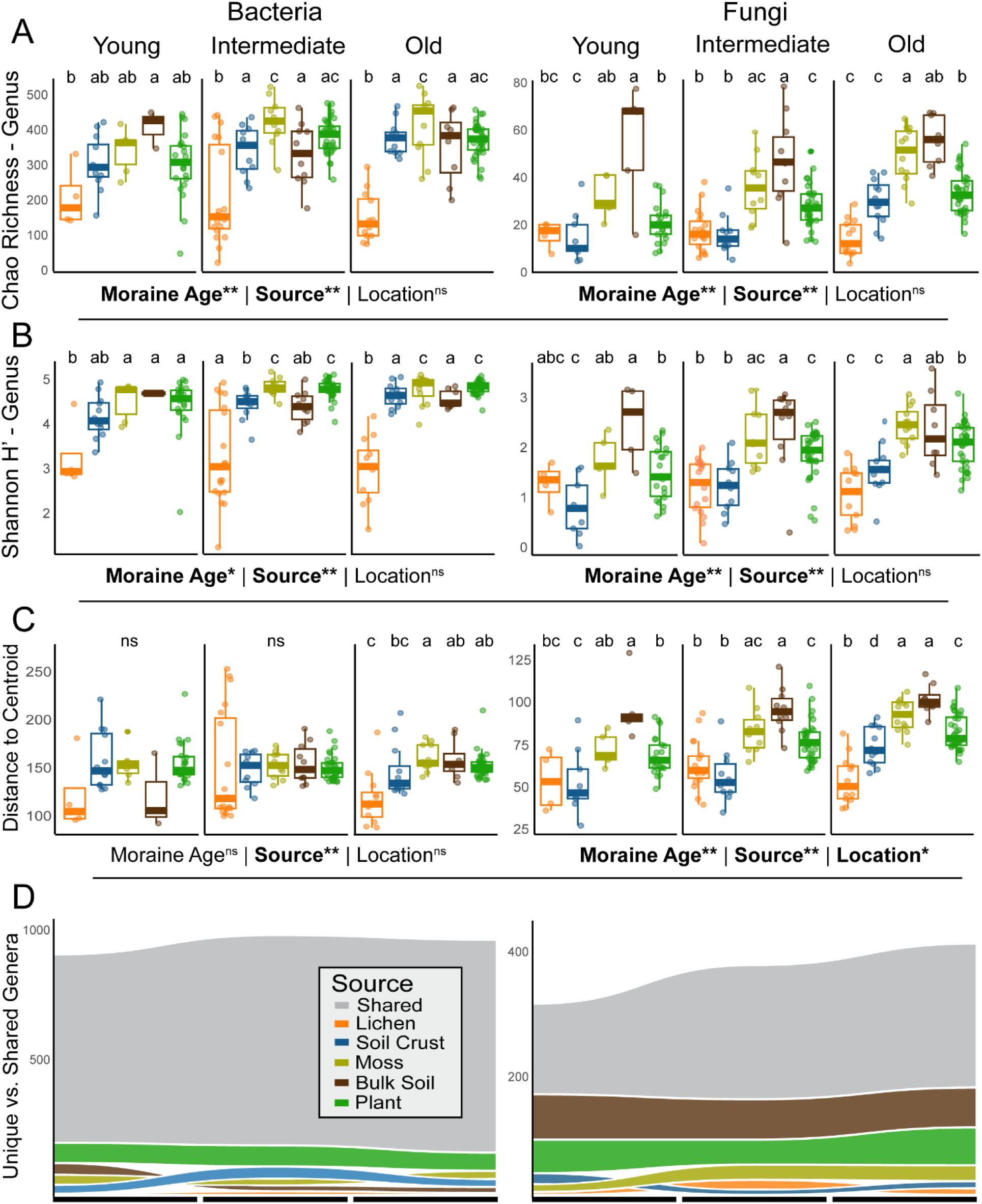
Boxplots (median and quartiles) of bacterial and fungal Chao richness (A), Shannon-Weiner (H’) diversity index (B), and beta-dispersion (C) at the genus level for each respective source across three categorized moraine ages (young, intermediate, old). Linear mixed-effect models were performed to test the effect of moraine age, source, and location with individual plots as a random factor. Bump plots (D) show the number of unique vs. shared genera found in each source across the categorized moraine ages (Young, Intermediate, Old).

### 3.4 Bacterial and fungal β-diversity

Community composition of both bacterial and fungal taxa followed similar trends, with biotic source, moraine age, and location all significantly influencing variation, as shown by PERMANOVA analyses (Figure 2A). Biotic source had the strongest effect, explaining 20.9% of variation in bacterial communities and 10.0% in fungal communities. Among bacteria, moraine age accounted for more variation (2.1%) than location (1.4%), whereas in fungal communities, location explained slightly more variation (3.2%) than moraine age (2.8%). When analyzed within each biotic source, bacterial community composition was significantly influenced by moraine age in moss and plant samples, and by location in soil crust, bulk soil, and plant samples. For fungal communities, moraine age significantly affected soil crust, moss, and plant samples, while location influenced communities in lichens, mosses, bulk soil, and plants (Supplementary Table 2). Beta-dispersion linear mixed-effect models showed source identity had the most significant effect upon bacterial communities. In contrast, fungal beta-dispersion was more dynamic, being affected by source identity and location, while generally increasing in older moraines (Figure 2C).

### 3.5. Plot parameters, source exclusion, and microbial co-occurrence patterns

Hierarchical RDAs revealed that plot cover measurements had the strongest influence on global network parameters. For both bacterial and fungal metacommunities, axis 1 of the cover-based PCA was the only significant explanatory variable for global network structure (p = 0.036 for bacteria; p = 0.002 for fungi; Figure 4A, B). In both cases, this axis primarily reflected a gradient of increasing plant cover and decreasing stone cover.

In the bacterial RDA, increasing plant cover also aligned with axis 2 of the environmental PCA, which represented increasing moraine age. Conversely, higher stone cover was associated with increasing coarse-grained soil texture. In total, the six axes from the PCAs constrained 25.7% of the variation. Average node dissimilarity, edge connectance, and vertex connectance were positively correlated with higher stone cover in younger moraine plots, indicating that bacterial metacommunities in these environments were highly interconnected. Although higher node dissimilarity may suggest that individual nodes (taxa) differed substantially in their associations. In contrast, older moraines with greater plant cover exhibited higher values of modularity, average path length, natural connectance, and perturbation effect profile, suggesting a more structured and compartmentalized metacommunity, where taxa form tighter, more internally connected subgroups. These patterns were further supported by source exclusion network analyses. When plant rhizosphere communities were excluded, edge connectance increased, resembling the structure observed in younger moraine plots (Figure 4C). Additionally, removing lichen-associated bacterial communities led to significant reductions in natural connectance and perturbation effect profile values (Supplementary Table 3). Collectively, these findings indicate that bacterial co-occurrence patterns are strongly influenced by the presence, distribution, and abundance of hosts and sources across the glacier moraine chronosequence.

Fungal metacommunities exhibited trends similar to those observed in bacterial communities, with the primary RDA axis reflecting moraine age and associated changes in soil properties. Together, the RDA axes explained 30.5% of the total variation in network structure (Figure 3B). Unlike the bacterial RDA, average node dissimilarity was the only network parameter strongly associated with younger moraines characterized by higher stone cover. This suggests that fungal metacommunities in early-successional environments are more fragmented and disconnected. In contrast, older moraines with greater plant cover and enriched soil nutrients were associated with increased values of modularity, natural connectance, perturbation effect profile (PEP), edge connectance, and vertex connectance, indicating that late-successional fungal metacommunities are more structured, with distinct subgroups that are also better integrated within the broader network. Excluding plant rhizosphere samples led to an increase in clustering coefficient and average path length (Figure 4D; Supplementary Table 4), suggesting that plant-associated fungal communities contribute to the formation of distinct subgroups and include taxa that serve as critical connectors across the metacommunity. Additionally, the removal of plant fungal communities resulted in a decrease in PEP values, whereas inclusion of all sources yielded the highest PEP values. This indicates that plant-associated fungi play a key role in maintaining metacommunity resilience, although contributions from all sources are important for overall network stability.

**Figure 3.**
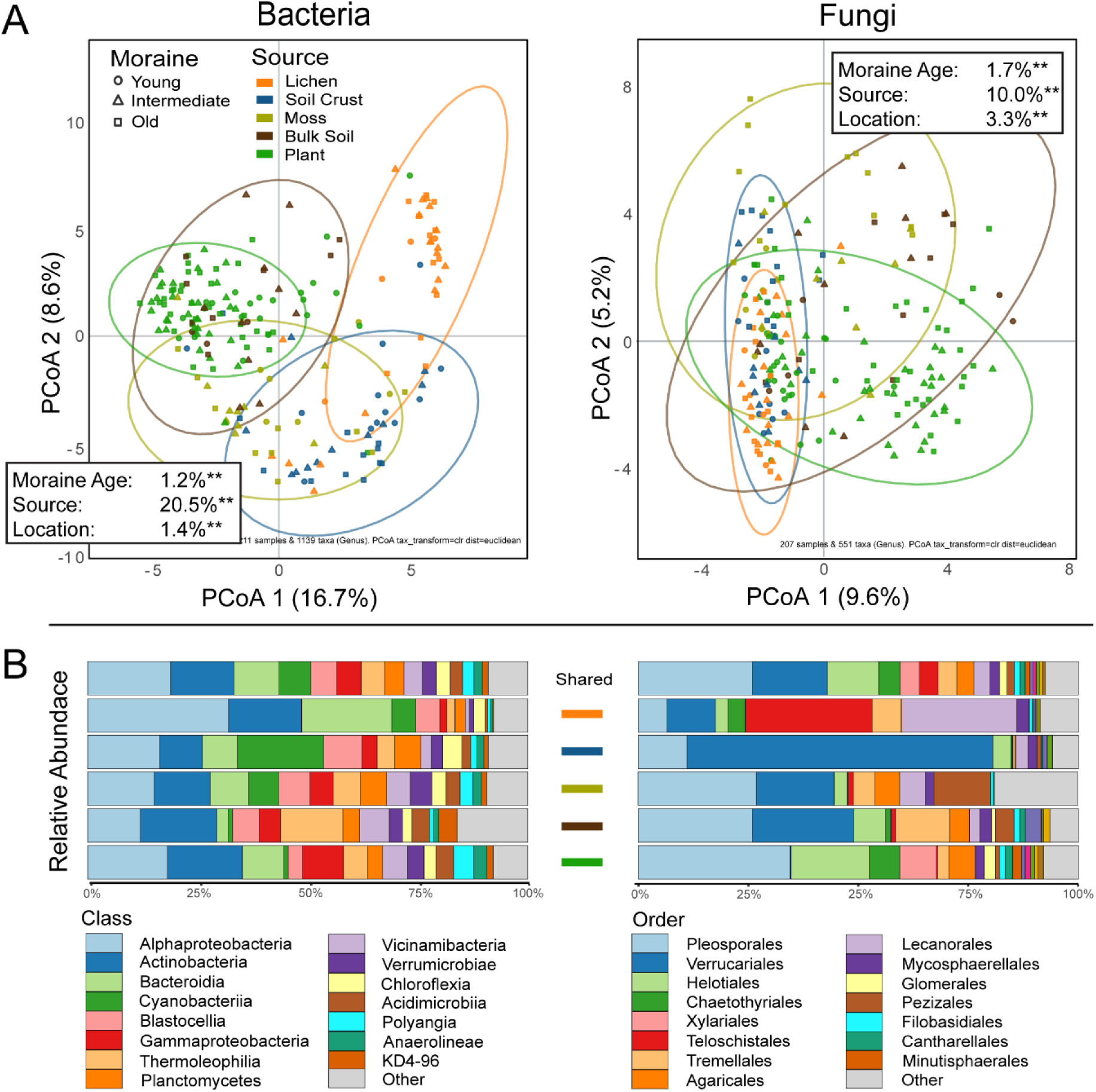
Principal Coordinates Analysis (PCoA) of bacterial (n = 211) and fungal (n = 207) genus-level composition, grouped by source and moraine age (A). Data was centered log-ratio (CLR) transformed and analyzed using Euclidean distance. Ellipses represent 95% confidence intervals for each source. PERMANOVA analyses were performed using the same transformation and distance calculations with individual plots as a random factor. Compositional bar plots (B) show the pooled relative abundance of higher taxonomic classifications for each respective source and shared genera across all sources.

**Figure 4.**
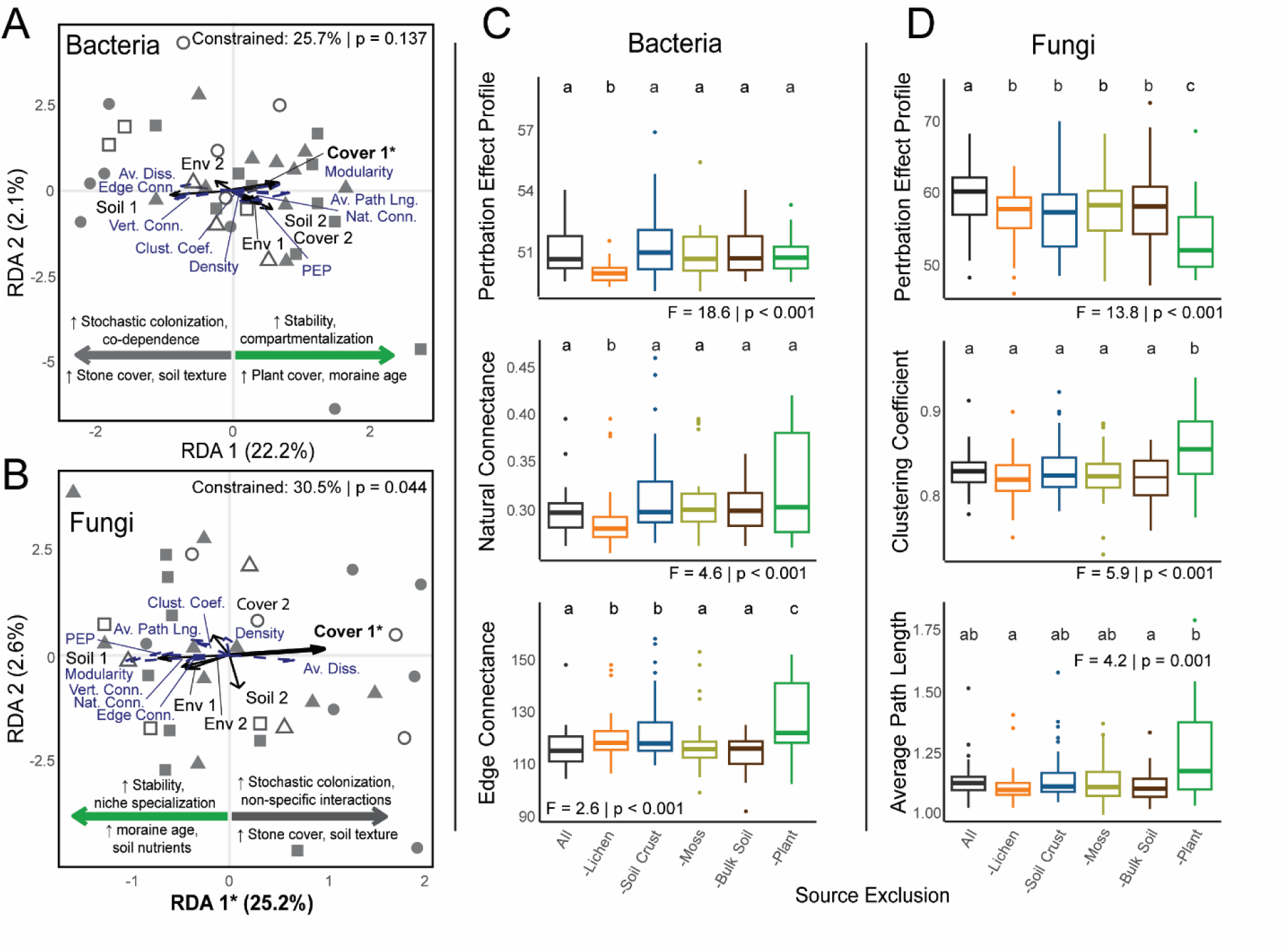
Hierarchical redundancy analysis (RDA) showing the influence of environmental variables (Env), plot cover measurements (Cover), and soil properties on RMS-scaled global network parameters of bacterial (A) and fungal (B) communities. RDA was performed on the first and second axes extracted from separate PCAs for each variable group, conditioned by sampling location. Boxplots of bacterial (C) and fungal (D) global network parameters assessed under separate source exclusion treatments representing median and quartiles. Data were analyzed using a non-parametric aligned rank transformation (ART) ANOVA with *plot* as a random effect. Pairwise comparisons of estimated marginal means were conducted post hoc, with p-values adjusted using the false discovery rate (FDR) correction.

### 3.6. Environmental Drivers of Bacterial and Fungal Community Congruence

Among the Procrustes analyses of all samples and individual sources, significant congruence between bacterial and fungal communities was detected only in moss-associated samples (Table 1). Analysis of Procrustes residuals revealed significant correlations with environmental factors: bulk soil communities showed a negative relationship with increasing distance from the glacier tongue (coef = –0.52; p = 0.016), while plant-associated microbial communities exhibited a positive correlation with the Snow 1976 variable (coef = 0.32; p = 0.015). Since higher Procrustes residuals indicate greater decoupling between bacterial and fungal communities, these results suggest that bacterial–fungal congruence in bulk soil increases with moraine age, whereas in plant-associated communities, congruence decreases in areas with prolonged snow cover. Marginally significant relationships were also observed between residuals and environmental variables, including soil crust communities and aspect (sine) (coef = –0.32; p = 0.078), bulk soil and elevation (coef = –0.40; p = 0.075), bulk soil and Snow 1976 (coef = 0.43; p = 0.054), and plant communities and distance from the glacier tongue (coef = –0.24; p = 0.083). These findings indicate that bacterial–fungal community congruence is highly context-dependent, with mosses showing the strongest alignment, while factors such as moraine age, snow persistence, and topographic features strongly influence the degree of bacterial–fungal coupling across sources.

**Table 1.** Procrustes analysis results assessing the congruence between bacterial and fungal community structures across all samples and within each source. The table reports Procrustes R values and corresponding p-values. Input ordinations were generated via redundancy analysis (RDA), constrained by Location. To identify environmental factors associated with community incongruence, linear mixed-effect models were applied using environmental variables as predictors of Procrustes residuals, with Plot ID included as a random effect (* p-value < 0.05).

|  |  | Sources |  |  |  |  |  |
| --- | --- | --- | --- | --- | --- | --- | --- |
|  |  | All | Lichen | Soil Crust | Moss | Bulk Soil | Plant |
| Procrustes | Num. samples | 201 | 34 | 31 | 26 | 21 | 89 |
|  | R | 0.08 | 0.20 | 0.22 | 0.40 | 0.31 | 0.18 |
|  | P-value | 0.44 | 0.50 | 0.41 | **0.04** | 0.35 | 0.12 |
| Linear Mixed-Effect Models | Distance from Glacier | 0.06 | 0.12 | -0.06 | 0.23 | **-0.52\*** | -0.25 |
|  | Elevation | -0.09 | -0.15 | -0.08 | 0.05 | -0.40 | -0.14 |
|  | TPI | 0.04 | 0 | -0.15 | -0.08 | 0.36 | 0 |
|  | Aspect (Cosine) | 0 | -0.20 | 0 | 0.07 | -0.35 | -0.02 |
|  | Aspect (Sine) | -0.08 | -0.25 | -0.32 | -0.01 | -0.12 | -0.02 |
|  | Slope | 0.09 | -0.12 | -0.09 | -0.01 | -0.04 | 0.04 |
|  | Snow 1976 | -0.04 | 0.15 | -0.06 | 0.15 | 0.43 | **0.32\*** |

### 3.7. Keystone bacterial and fungal genera

Among the ten keystone genera identified using the dual-criteria approach for bacteria and fungi, nine of the top ten bacterial genera were associated with plant rhizosphere samples, with only one originating from bulk soil (Table 2). In contrast, fungal keystone genera were more evenly distributed, with six identified from bulk soil and four from plant rhizosphere samples (Table 3). Bacterial keystone genera were present in 24 to 46 total networks in which they had an eigenvalue > 0.95, while fungal genera ranged from 13 to 41 such networks in which they had an eigenvalue > 0.95. Maximum relative abundances for bacterial keystone genera ranged from 5% to 1.1%, whereas fungal genera exhibited a much broader range—from 54% down to 1.1%. Three bacterial keystone genera (Roseiflexaceae *incertae sedis* gen.1, KD4-96 *incertae sedis* gen.1, and *Gaiella*) were significantly negatively correlated with the Snow 1976 variable, suggesting that their abundance increases in plant rhizospheres where bacterial and fungal communities become more tightly coupled. Similarly, three fungal keystone genera, *Polyblastia*, *Xanthoria*, and *Coprinus*, showed increased abundance with greater distance from the glacier tongue in bulk soil, coinciding with stronger bacterial–fungal congruence in those sites.

**Table 2.** Keystone bacterial genera identified across glacier forefields in the Ladakh region of northwest India. Genera were selected based on the following criteria: maximum relative abundance > 0.01, significant correlation with Procrustes residuals (Spearman rank p < 0.01), and high network centrality (eigenvector centrality > 0.95). Taxa were ranked according to the number of networks in which they were identified as keystone species. Bold indicates significance for residuals and environmental variables.

| Rank | Genus | Max (%) | Total Networks | Source | Spearman Rank Test |  |  |  |  |  |
| --- | --- | --- | --- | --- | --- | --- | --- | --- | --- | --- |
|  |  |  |  |  | Procrustes Residuals |  | Snow 1976 |  | Distance From Glacier |  |
|  |  |  |  |  | R<sup>2</sup> | p | R<sup>2</sup> | p | R<sup>2</sup> | p |
| 1 | **Roseiflexaceae inc. sed. gen.1** | **5.0%** | **46** | **Plant** | **-0.337** | **0.001** | **-0.244** | **0.021** | - | - |
| 2 | Microscillaceae inc. sed. gen.1 | 3.9% | 37 | Plant | -0.301 | 0.004 | -0.043 | 0.692 | - | - |
| 3 | **KD4-96 inc. sed. gen.1** | **4.2%** | **35** | **Plant** | **-0.300** | **0.004** | **-0.282** | **0.007** | - | - |
| 4 | Ilumatobacteraceae inc. sed. gen.1 | 1.4% | 34 | Bulk Soil | -0.470 | 0.033 | - | - | -0.090 | 0.699 |
| 5 | **Gaiella** | **2.4%** | **26** | **Plant** | **-0.338** | **0.001** | **-0.240** | **0.024** | - | - |
| 6 | SBR1031 inc. sed. gen.1 | 1.7% | 26 | Plant | -0.306 | 0.004 | -0.144 | 0.177 | - | - |
| 7 | CL500-29 inc. sed. gen.1 | 2.4% | 25 | Plant | -0.231 | 0.030 | 0.049 | 0.649 | - | - |
| 8 | *Mesorhizobium* | 1.6% | 24 | Plant | -0.345 | <0.001 | -0.208 | 0.050 | - | - |
| 9 | SJA-28 inc. sed. gen.1 | 1.1% | 24 | Plant | -0.234 | 0.027 | -0.148 | 0.166 | - | - |
| 10 | C0119 inc. sed. gen.1 | 1.2% | 24 | Plant | -0.355 | <0.001 | -0.069 | 0.518 | - | - |

**Table 3.** Keystone fungal genera identified across glacier forefields in the Ladakh region of northwest India. Genera were selected based on the following criteria: maximum relative abundance > 0.01, significant correlation with Procrustes residuals (Spearman rank p < 0.01), and high network centrality (eigenvector centrality > 0.95). Taxa were ranked according to the number of networks in which they were identified as keystone species. Bold indicates significance for residuals and environmental variables.

| Rank | Genus | Max (%) | Total Networks | Source | Spearman Rank Test |  |  |  |  |  |
| --- | --- | --- | --- | --- | --- | --- | --- | --- | --- | --- |
|  |  |  |  |  | Procrustes Residuals |  | Snow 1976 |  | Distance From Glacier |  |
|  |  |  |  |  | R<sup>2</sup> | p | R<sup>2</sup> | p | R<sup>2</sup> | p |
| 1 | ***Polyblastia*** | **27.8%** | **41** | **Bulk Soil** | **-0.471** | **0.031** | - | - | **0.567** | **0.007** |
| 2 | *Debaryomyces* | 6.5% | 33 | Bulk Soil | -0.440 | 0.046 | - | - | 0.428 | 0.053 |
| 3 | ***Xanthoria*** | **54.2%** | **29** | **Bulk Soil** | **-0.550** | **0.010** | - | - | **0.483** | **0.027** |
| 4 | ***Coprinus*** | **1.1%** | **28** | **Bulk Soil** | **-0.435** | **0.049** | - | - | **0.520** | **0.016** |
| 5 | *Tomentella* | 1.1% | 25 | Bulk Soil | -0.457 | 0.037 | - | - | 0.423 | 0.056 |
| 6 | *Cladosporium* | 12% | 18 | Plant | -0.231 | 0.029 | -0.175 | 0.102 | - | - |
| 7 | *Extremus* | 1.1% | 18 | Bulk Soil | -0.511 | 0.018 | - | - | 0.244 | 0.286 |
| 8 | Microsphaeropsidaceae<br>*inc. sed. gen.1* | 1.9% | 15 | Plant | 0.239 | 0.024 | 0.035 | 0.746 | - | - |
| 9 | *Laetisaria* | 38.8% | 14 | Plant | -0.223 | 0.036 | -0.005 | 0.966 | - | - |
| 10 | *Cistella* | 34.4% | 13 | Plant | -0.209 | 0.049 | -0.109 | 0.308 | - | - |

## 4. Discussion

As the initial colonizers and foundation of biogeochemical cycling in glacier forefields, understanding the influence of unique sources upon microbial community assembly is paramount to preserving biodiversity and maintaining ecosystem stability over long-term succession. Bacterial and fungal communities are shaped by a multitude of interacting abiotic and biotic factors, as changing climate regimes and newly exposed glacial terrain lead to the rapid colonization by soil crusts, cryptogams, and vascular plants. This study extends the existing knowledge by addressing the importance of these hosts and abiotic sources (lichens, BSCs, mosses, bulk soil, and plants) in shaping microbial diversity and investigating their role in microbial metacommunity assembly processes on a small scale and across wide environmental gradients of soil development, topography, and changing cover measurements at four glacier forefields in the high desert region of Ladakh, India (northwestern Himalaya). Using a multidisciplinary chronosequence approach, integrated with GIS-based terrain analysis, molecular sequencing, and classical transect sampling, we confirmed that moraine age is the dominant driver of abiotic soil properties and vegetation cover, which subsequently influences bacterial and fungal diversity and metacommunity assembly (Figure 5). As hypothesized, microbial alpha- and beta-diversity varied across sources, with lichens consistently hosting the lowest diversity and plants and mosses supporting richer communities in older moraines. Furthermore, we demonstrate that topographic depressions which were once regularly snow-covered are partially delayed in their succession and affect the congruence between bacterial and fungal communities. Co-occurrence network analyses revealed a shift from low-modularity communities in young, stone-covered plots to more structured, compartmentalized networks in older, plant-covered environments. Finally, using a dual-criteria approach, we identified keystone genera from bacterial and fungal communities, some of which corresponded with environmental gradients and inter-kingdom congruence.

**Figure 5.**
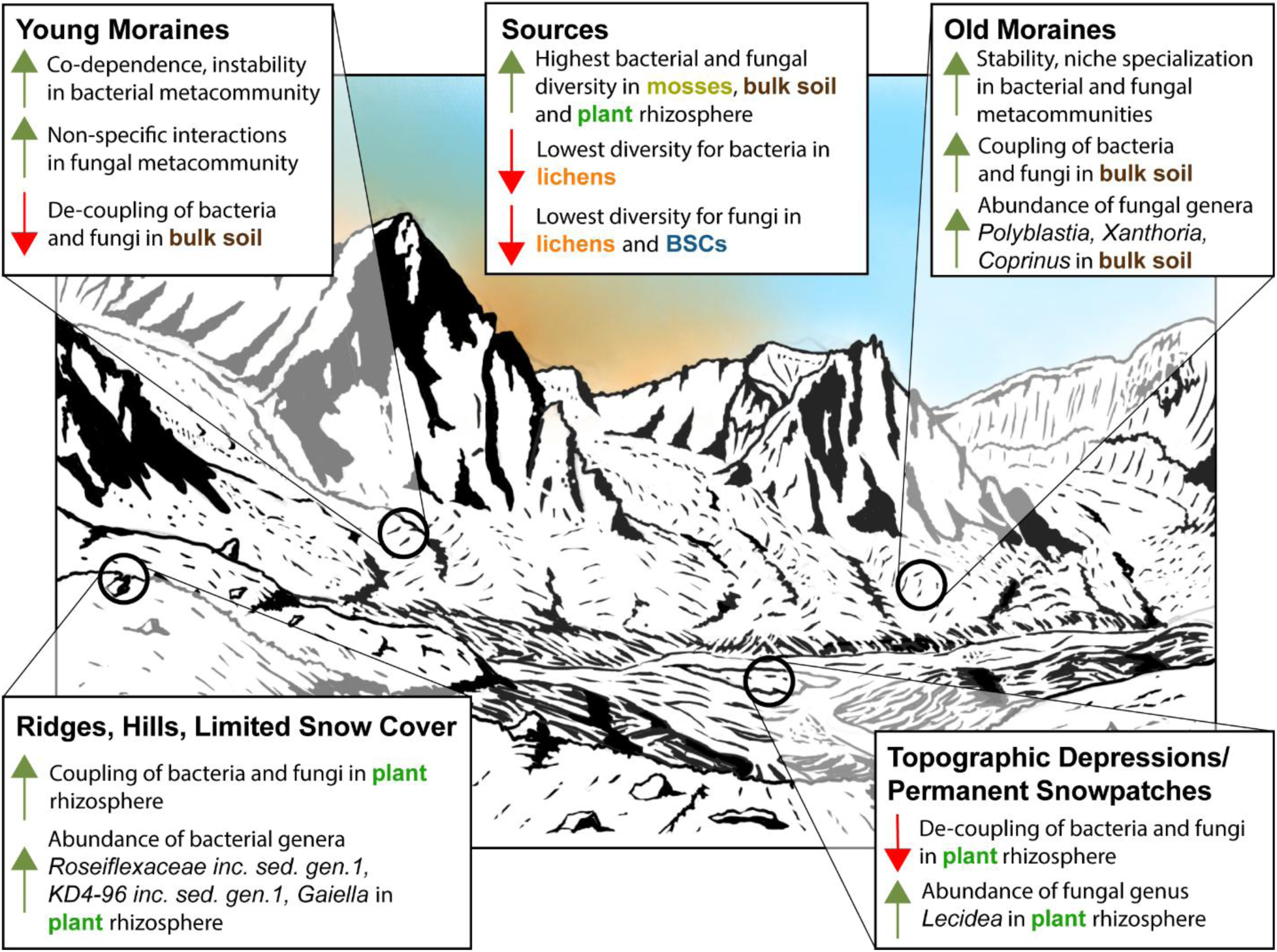
Conceptual diagram illustrating microbial metacommunity assembly processes across glacier forefields in Ladakh, India. The figure synthesizes key findings from bacterial and fungal co-occurrence network analyses, the relationships between environmental gradients and Procrustes residuals within specific sources, and the identification of keystone fungal genera responsive to environmental factors. Illustration drawn by ATR.

### 4.1. Microbial diversity across sources

The distinct microbial communities associated with colonizing macroorganisms and the bulk soil reflect a combination of successional status, microhabitat differentiation, and the selective pressures exerted by the host. Lichens, often among the earliest colonizers of post-glacial environments, harbor a notably specific holobiome compared to other sources, maintaining low microbial diversity and beta-dispersion, while hosting few unique taxa throughout the chronosequence (Figure 2). Growing primarily on exposed rock surfaces, lichens endure extreme conditions, including intense UV radiation, desiccation, and nutrient limitation. These harsh environmental factors are reflected in the composition of their associated bacterial communities, which are often dominated by resilient taxa such as Alphaproteobacteria and Actinobacteria (Touchette et al. 2023, Tagirdzhanova et al. 2025). Yet, there is growing evidence that the microbiota is metabolically linked to the lichen thallus, providing vitamins and micronutrients while also mitigating pathogens and environmental stressors (Cernava et al. 2017). Furthermore, the consistent observation of *Acidiphilium* and *Sphingomonas* within the lichen microbiome in our study and others (Li et al. 2023b, Tagirdzhanova et al. 2025), shows these relationships are persistent as they approach the upper limits of their elevational distribution. Amongst the dominant fungal taxa within the lichen thallus, *Knufia* and *Dioszegia* appear to be extremophilic commensals (Isola et al. 2016, Chander et al. 2024), with the later having also been isolated from the lichens in the High Arctic (Zhang et al. 2015). These findings suggest that the lichen holobiome consists of a mix of commensal and potentially mutualistic relationships, although the precise symbiotic status of many of these interactions remains unresolved.

Biological soil crusts, also among the early colonizers in a glacier forefield, exhibited a more dynamic successional trajectory in their microbial communities, potentially reflecting their deeper integration with the broader ecosystem (Schmidt et al. 2025). These findings corroborate our previous work, which documented a progressive shift in dominant cyanobacterial taxa within BSCs at the same study locations (Ruka et al., 2023). While overall BSC microbial diversity was often lower than that observed in other sources (apart from lichens), as well as the number of unique taxa hosted by them, changes in alpha-diversity metrics and fungal beta-diversity across moraine ages indicate that BSC succession is not limited to cyanobacteria alone (Zhou et al. 2023). *Sphingomonas* is a common constituent of BSC bacterial communities (Zhang et al. 2016, Zhou et al. 2023), while *Blastocatella* appears to be a taxon more related to the arid, high pH conditions characteristic of the Ladakh region (Foesel et al. 2013). Fungal communities within BSCs were dominated by members of the Verrucariales, aligning with other studies reporting high relative abundances of Ascomycota in arid BSCs (Bates et al. 2012, Borchhardt et al. 2019, Liu et al. 2024). These fungal taxa, often exhibiting a lichenized growth, are commonly found in BSCs in arid regions and contribute to soil aggregation and enhanced desiccation resistance (Zhang et al. 2017, Guang et al. 2025). As BSCs shift from cyanobacteria-dominated to more complex communities, the stabilizing role of fungi highlights the need for further study of their ecological function in high-altitude, arid environments.

Within a glacier forefield, mosses exist in a plethora of microniches, from growing in rock cracks, within the BSC matrix, to the base of plants, often associated with areas of higher moisture. Despite mosses lacking true roots, there is growing evidence of mosses harboring specific microbial communities (Alvarenga and Rousk 2022, Ramakrishnan et al. 2025), even across their entire lifecycle (Bragina et al. 2012). Our study adds to this knowledge by demonstrating that moss-associated communities become more diverse over successional time, with both bacterial and fungal communities significantly influenced by moraine age. Most importantly, mosses were the only source to show a significant coupling between bacterial and fungal communities, revealing a previously unrecognized association within the moss microbiome. While the dominant bacterial taxa resembled those in soil crusts, cyanobacteria were less abundant, suggesting that mosses supplant the carbon source and subsequently shape the microbial community through higher C:N ratio tissue and secondary metabolites, such as antimicrobial products (Klarenberg et al. 2020, Valeeva et al. 2022). Similar fungal taxa, such as Verrucariales and Pleosporales, were particularly abundant in the moss holobiome, indicating ecological overlap with soil crusts. However, the high prevalence of *Lamprospora* (Pezizales), a common moss parasite, within moss fungal communities, demonstrates how these ecological interactions extend from milder zones to some of the highest elevations ever studied (Vega et al. 2019). Yet, ongoing climate change threatens moss prevalence within glacier forefields through moisture loss and habitat shifts (Wookey et al. 2009), potentially disrupting the added stability mosses provide to microbial communities, even as their high microbial diversity and consistent bacterial-fungal congruence suggest they serve as a critical microbial reservoir shaped by moss species identity and interdependent ecological interactions.

Bulk soil microbial communities likely function as a reservoir for the broader metacommunity within glacier forefields (Chen et al. 2023), demonstrated by high bacterial and fungal alpha diversity. However, unsuccessful sequencing attempts from young moraine bulk soils suggest there is extremely low microbial biomass in early successional stages (Jiang et al. 2018, Feng et al. 2023), where other sources may provide essential nutrient inputs that promote microbial growth and community differentiation. The high abundance of *Solirubrobacter* (Thermoleophilia), a potentially mobile, UV-resistant bacterium often associated with plant roots, further supports the idea that bulk soil communities are shaped by microbial recruitment from nearby plants (Ruka et al., 2023). Fungal communities in bulk soil included many generalist taxa, but demonstrated increased abundance of Tremellales, particularly *Vishniacozyma*, a cold-adapted yeast isolated from glacial sediments in the Canadian High Arctic, capable of growing at sub-zero temperatures under nutrient-poor conditions (Tsuji et al. 2019). Thus, these findings suggest that bulk soil communities consist largely of generalists shared with other sources, while also harboring cold-tolerant specialists adapted to extreme forefield environments.

The selective capabilities of plants within the rhizosphere, through root exudates and decaying plant material, have established them as a recognized “hotspot” of microbial diversity (Kuzyakov and Blagodatskaya 2015). Our study confirms that plant-associated communities generally exhibit higher microbial diversity than other sources; however, this selective environment may also constrain the overall number of taxa and evenness compared to mosses or bulk soil. Rhizosphere microbial diversity was clearly influenced by moraine age (Ciccazzo et al. 2014), particularly among fungi, and contributed a number of unique taxa across the chronosequence, although we did not directly investigate whether dispersed plant seeds introduce their own microbiota. We observed notable increases in fungal orders such as Pleosporales and Helotiales, consistent with findings from another rhizosphere study in Ladakh (Kotilínek et al. 2017, Hiiesalu et al. 2023, Hussain et al. 2024). While these groups commonly include saprotrophs, facilitating the decomposition of plant detritus within the rhizosphere, they also consist of fungal symbionts, such as ericoid mycorrhiza and dark-septate endophytes which are known to confer stress resistance in cold habitats (Malicka et al. 2022). Some taxa, such as *Comoclathris* and *Alternaria,* were widely distributed across multiple sources, whereas others engaged more exclusively with their hosts through endophytic symbioses (*Nothodactylaria, Exophiala*) (Maciá-Vicente et al. 2016, Li et al. 2024). Among bacterial communities, an increase in Gammaproteobacteria was observed, including *Steroidobacter*, a genus known to produce brassinosteroids, plant hormones that promote cell growth and enhance stress tolerance protection (Fahrbach et al. 2008). Furthermore, *Steroidobacter* has been reported in the rhizosphere community of *Welwitschia* plants, suggesting that alpine plants in high-elevation glacier forefields may utilize similar microbial strategies to cope with environmental extremes (Valverde et al. 2016). Together, these results underscore the rhizosphere’s critical role in selectively shaping microbial communities, harboring both ubiquitous species and rhizosphere-specific taxa.

### 4.2. Environmental factors affecting inter-kingdom congruence

Topographic heterogeneity in glacier forefields does not act independently from moraine age and soil development, making it difficult to disentangle these overlapping effects (Yamagata 2022, Masumoto et al. 2023). While previous studies have shown the influence of topography on single taxonomic groups (Solon et al. 2021, Masumoto et al. 2023, Ruka et al. 2023), our attempt to assess its effect on bacterial–fungal interactions found limited results. The higher inter-kingdom congruence observed in bulk soils of older moraines may partly stem from the lack of successfully sequenced samples in younger moraines. However, it may also reflect the relatively stable conditions in later successional stages, characterized by increased OM, TN, and micronutrients, as well as greater plant cover that buffers soil temperature and moisture extremes (Philippot et al. 2011, Donhauser and Frey 2018). Interestingly, low-lying areas that remained snow-covered during the summer of 1976 in our study area exhibited reduced bacterial–fungal congruence in plant rhizospheres. This variable is multi-faceted and heavily influenced by moraine age, as sites closer to glaciers tend to be colder, have longer snow cover, and shorter growing seasons (Magnusson et al. 2010). Such depressions are also more likely to be snow-covered during mid-season precipitation events, buffering microbial communities from extreme temperatures but limiting plant-driven carbon inputs (Jandova et al., 2025). These areas may lag behind in successional development, having only recently become exposed compared to nearby ridges and hills that are more frequently snow-free. Thus, the intersection between changing climatic regimes and its effect upon multi-trophic interactions suggests a “soil legacy” effect within glacial chronosequences which is not observed during a single field campaign (Dacal et al. 2022, Li et al. 2023a). Although this study did not explicitly assess these dynamics, our findings suggest that topographically mediated snow cover may be an overlooked factor in chronosequence studies, potentially affecting microbial interactions via altered plant performance. Snow-free areas tend to support advanced vegetative communities, with earlier phenology and higher recruitment (Jandova et al., 2025, Chen et al. 2025). In contrast, prolonged snow cover may lead plants to shift biomass allocation toward stress tolerance rather than investing in microbial relationships (Zhang et al. 2023). Ecologically, higher inter-kingdom congruence may indicate tighter symbiotic associations and more efficient nutrient cycling, reflecting coordinated responses among microbial kingdoms that enhance nutrient availability and plant productivity (Zhao et al. 2017, Zhou et al. 2022). Conversely, decreased congruence could suggest weaker or more competitive interactions, potentially associated with pathogenic dynamics or reduced host selectivity in stressful environments. In summary, while separating environmental influences from moraine age remains a challenge, there are likely underlying relationships which influence inter-kingdom relationships within hosts and sources due to physiological stress and shifts in climatic patterns.

### 4.3. Metacommunity assembly processes across a glacier chronosequence

Microbial community structure is shaped by a multitude of factors including positive and negative interactions, environmental heterogeneity, stochastic dispersal and deterministic selection by higher trophic levels. In glacier forefields, these factors are further constrained by an overarching temporal gradient (Matthews 1992, Bradley et al. 2014). Using co-occurrence network analyses, we found that bacterial and fungal community structures were primarily influenced by increasing plant cover in older moraines vs. stone cover in younger moraines. Notably, both communities exhibited greater average node dissimilarity in younger moraines, indicating that taxa are assembled more stochastically at early successional stages. However, bacterial communities in these younger sites also showed increased edge connectivity, suggesting that despite their stochastic assembly, they form increasingly co-dependent networks, for example, in soil crusts where cyanobacteria contribute nitrogen fixation and carbon assimilation, while heterotrophic bacteria support decomposition and mineralization (Belnap et al. 2003, Schmidt et al. 2025). In contrast, although fungi also inhabit these early successional environments, their interactions with the community appear to be less defined at this stage. However, in later successional stages, both bacterial and fungal networks demonstrated increased modularity, average path length, and natural connectance, indicative of more defined interactions and greater structural stability (Newman 2006). This suggests that as the metacommunity diversifies, functional redundancy within modules may enhance resilience to disturbance events. These patterns partly align with findings from other alpine studies; for example, early successional microbial communities in the Alps showed similar network trends (Mandolini et al. 2025), although Vimercati et al. (2022) observed decreasing modularity in bacterial communities over time. A key difference, however, is that our study incorporated multiple sources into the network analyses, while previous studies focused solely on bulk soils.

Our dual criteria approach to identifying keystone taxa, focused on bulk soil and rhizosphere communities, revealed that most bacterial keystone taxa were associated with the plant rhizosphere and became more abundant in environments with higher inter-kingdom congruence. This suggests that these bacterial keystones taxa may thrive under optimal plant conditions and through synergistic interactions with fungal counterparts (Cui et al. 2025, Cheng and Ma 2025). Many of these taxa remain uncultured, indicating that they may be rare specialists with strict nutrient or symbiotic requirements (Vartoukian et al. 2010). Keystone fungal taxa were predominantly identified in bulk soils and mostly increased in abundance when inter-kingdom congruence increased. In older moraines, lichenized fungi were surprisingly prevalent in bulk soils, potentially due to their role in soil aggregation and plant facilitation (Santiago et al. 2018), though the possibility of detecting spores or extracellular DNA cannot be excluded. Meanwhile other taxa, such as *Coprinus* and *Cistella*, fulfilled more typical roles, contributing to soil organic carbon transformation (Zhang et al. 2024) or forming ericoid mycorrhizal associations (Koizumi and Nara 2017, Guo et al. 2024). Furthermore, our source exclusion analyses showed that the inclusion of plant rhizosphere communities effectively reshape microbial networks and increase interactions between microbial taxa (Cheng and Ma 2025). Together, these findings point to the pivotal role of plants not only in promoting keystone taxa but also in fostering complex inter-kingdom relationships that underpin microbial metacommunity assembly in glacier forefields.

## 5. Conclusion

Macroorganisms and bulk soil, acting as hosts and reservoirs of microbial diversity, create numerous microniches that strongly influence microbial community structure while experiencing a suite of environmental stressors, a factor especially critical in the heterogeneous landscape of a glacier forefield. Our findings, based on microbial succession in the high-desert Himalaya, reveal that unique sources, such as lichens, soil crusts, mosses, vascular plants, and bulk soil, harbor distinct communities that shift with moraine age. As these organisms are also early colonizers in other extreme environments, these results have broader relevance to alpine and polar ecosystems (Favero-Longo et al. 2012, Anderson et al. 2025). While separating environmental effects from moraine age remains difficult, our analysis showed that low-lying, geomorphological depressions with prolonged snow cover disrupted bacterial-fungal congruence in plant rhizospheres. In contrast, inter-kingdom congruence increased in bulk soil over time and remained consistently high in mosses, pointing to the selective and stabilizing role of mosses in microbial interactions. Vegetation cover, in conjunction with moraine age and rising nutrient availability, was a primary driver of microbial metacommunity structure, with younger sites showing unstable networks with simple structure and older sites developing modular, resilient networks with functional redundancy. Plants hosted the most keystone taxa, especially under conditions of strong inter-kingdom congruence, suggesting a link between plant performance and microbial network cohesion. Looking forward, this study supports a broader approach to studying microbial community assembly, while maintaining appropriate spatial scales, to investigate often overlooked sources of microbial diversity and the role of topography in shaping microbial and inter-kingdom interactions.

## Acknowledgements

ATR, RA and KŘ were supported by the Czech Science Foundation (GA ČR), grant numbers 21-04987S, 24-11954S and RVO 67985939, 60077344. JDa and IH were supported by Estonian Research Council grants PRG2584, PRG1789 and Center of Excellence “AgroCropFuture” and Estonian Academy of Sciences Estonia-Czech scientific cooperation project number EAS-EE-CZ-24-2.The funders had no role in study design, data collection and interpretation, or the decision to submit the work for publication. Also, we would like to thank Eva Petrová and Dana Švehlová for assisting in molecular methods and analyses.

## Open Research Statement

The short-read amplicon sequencing data will be deposited under the NCBI BioProject (made availability after acceptance). For reproducibility, reusability, and transparency, the scripts and data used in this study were deposited to GitHub (made available after acceptance). All functions and packages are cited within the paper without any associated novel code being produced for the results.

## Author Contributions

ATR, JDo and VL developed the experimental design. ATR, VL, SR performed the field work. KČ, KŘ, LV, JK performed the lab work. ATR performed the data analysis with assistance from IH, JDa and RA. ATR wrote the manuscript with significant contributions from KŘ and IH. TC and JDo were were consulted and advised interpretations of results. All coauthors read and approved the final version of this manuscript.

## 8. Supplementary Material

### 8.1 Appendix 1

Supplementary Methods and Results

#### 8.1.1 Moss and lichen identification

Mosses were identified at the University of South Bohemia in České Budějovice, Czechia, using a combination of classical morphological methods and molecular barcoding. This approach was necessary due to the limited and often inadequate taxonomic literature available for many moss groups in the region. DNA isolation, PCR amplification, and sequencing followed the protocols described in Kučera et al. (2019). The NaOH method was used for total genomic DNA extraction (Werner et al., 2002). Crude extracts were 10x diluted (chloroplast loci amplification) or 100x (ITS amplification) with 100 mM Tris-HCl (pH 8.3). For polymerase chain reactions, 0.5 μl DNA solution (10 μl final volume) was mixed with Plain PP MasterMix kit (Top-Bio, Vestec, Czech Republic) for normally amplifying samples and TP 2× Master Mix (Top-Bio) for amplification of ITS templates. Molecular identification was subsequently based on reference sequences from GenBank, targeting the nuclear ITS region, the plastid *trnL–trnF* intron and spacer, and/or the *rps4* gene.

Lichen identification was performed at Charles University in Prague, Czechia. Total genomic DNA was extracted directly from the lichen thallus using the cetyltrimethylammonium bromide (CTAB) protocol (Cubero et al., 1999). The fungal internal transcribed spacer (ITS) region (ITS1-5.8S-ITS2 rDNA) was amplified using the primers ITS1F (5’-CTT GGT CAT TTA GAG GAA GTA A-3’) (Gardes and Bruns, 1993) and ITS4 (5’-TCC TCC GCT TAT TGA TAT GC-3’) (White et al., 1990). Polymerase chain reactions (PCRs) were performed in 10 μl reaction volumes consisting of 6.5 μl Milli-Q water, 2 μl MyTaq reaction buffer, 0.1 μl MyTaq DNA polymerase (Bioline), 0.2 μl of each primer (25 mM), and 1 μl of template DNA. Successful PCR products were purified using AMPure XP SPRI paramagnetic beads (Beckman Coulter), following the manufacturer’s protocol. Sequencing was performed using the same ITS1F and ITS4 primers. The identity of the mycobiont was verified through sequence similarity searches using the BLAST tool (NCBI).

#### 8.1.2 Taxonomic description of bacterial and fungal communities within sources

The composition of bacterial communities varied across sources, particularly at higher taxonomic levels. Lichens were dominated by Alphaproteobacteria (*Acidiphilium*, *Sphingomonas*), Bacteroidiia (*Hymenobacter*, *Spirosoma*), and Actinobacteria (*Friedmanniella*). Soil crusts showed high abundances of Cyanobacteria (*Nostoc*, *Tychonema*, *Aliterella*), Blastocatellia (*Blastocatella*), and Alphaproteobacteria (*Sphingomonas*). Moss-associated communities were highly diverse, with dominant genera from Blastocatellia (*Blastocatella*, *RB41*), Thermoleophilia (*Solirubrobacter*), Cyanobacteria (*Nostoc*), and Alphaproteobacteria (*Sphingomonas*). Bulk soil communities were abundant in Thermoleophilia (*Solirubrobacter*, *Gaiella*) and Vicinamibacteria (unidentified genera), while Bacteroidiia were less abundant. Rhizosphere communities of plants had higher proportions of Gammaproteobacteria (*Steroidobacter*) and Polyangia (unidentified genera), along with persistent abundances of Alphaproteobacteria (*Sphingomonas*) and Actinobacteria (*Nocardioides*, *Actinoplanes*). Overall, these findings indicate broad taxonomic shifts among sources, yet certain genera, such as *Sphingomonas* and *Blastocatella*, remain consistently abundant across multiple sources.

Fungal communities exhibited strong specificity to sources, with substantial variation at the order level. In lichens, symbiotic fungi from the orders Teloschistales (*Xanthoria*) and Lecanorales (*Lecidella*, *Rhizoplaca*) were commonly abundant. Additionally, other fungal taxa often co-occurred with the primary symbionts, including *Knufia* (Chaetothyriales) and *Dioszegia* (Tremellales). Soil crust communities were primarily dominated by Verrucariales (*Endocarpon*, *Atla*, *Polyblastia*, *Verrucaria*), with notable contributions from Pleosporales (*Alternaria*) and Helotiales (*Tetracladium*). Moss-associated fungal communities were rich in Pleosporales (*Comoclathris*, *Phaeosphaeria*), Verrucariales (*Endocarpon*, *Agonimia*, *Polyblastia*, *Atla*), and Pezizales (*Lamprospora*). Bulk soil fungal communities also showed high abundances of Pleosporales (*Comoclathris*, *Alternaria*, *Cohesyomyces*, *Paraphoma*) and Verrucariales (*Endocarpon*), alongside elevated abundances of Tremellales (*Vishniacozyma*). In plant rhizosphere communities, dominant fungal taxa included Pleosporales (*Comoclathris*, *Alternaria*, *Darksidea*, *Paraphoma*), Helotiales (*Tetracladium*, *Cistella*), Chaetothyriales (*Exophiala*, *Cyphellophora*, *Fonsecaea*), and Xylariales (*Nothodactylaria*). As with bacterial communities, certain fungal taxa, such as *Endocarpon* and *Comoclathris*, were broadly distributed and consistently abundant across multiple sources, despite significant taxonomic shifts at the order level.

### 7.2 Appendix 2

#### Supplementary Tables and Figures

**Supplementary Table 1.** Identification and distribution of sequenced microbial community samples across moraine ages, locations, and sources.

| Source | Young |  |  | Intermediate |  |  | Old |  |  | Grand Total |
| --- | --- | --- | --- | --- | --- | --- | --- | --- | --- | --- |
|  | NUB | TSO | Total | NUB | TSO | Total | NUB | TSO | Total |  |
| **Bulk Soil** | 3 |  | 3 | 7 | 3 | 10 | 6 | 2 | 8 | 21 |
| **Soil Crust** | 9 | 3 | 12 | 8 | 3 | 11 | 9 | 3 | 12 | 35 |
| **Lichen** | 3 | 1 | 4 | 14 | 5 | 19 | 10 | 3 | 13 | 36 |
| *Xanthoria elegans* | 2 | 1 | 3 | 6 | 3 | 9 | 4 | 2 | 6 | 18 |
| *Gyalolechia bracteata/desertorum* |  |  |  | 4 |  | 4 |  |  |  | 4 |
| *Lecidea tessellata* |  |  |  |  |  |  | 2 |  | 2 | 1 |
| *Aspicilia* sp. |  |  |  |  |  |  | 1 |  | 1 | 1 |
| *Lecanora* sp. |  |  |  | 1 |  | 1 |  |  |  | 1 |
| *Lecidella stigmatea/patavina* |  |  |  |  |  |  | 1 |  | 1 | 1 |
| *Rhizoplaca melanophtalma* |  |  |  | 1 |  | 1 |  |  |  | 1 |
| Unidentified | 1 |  | 1 | 2 | 2 | 4 | 2 | 1 | 3 | 8 |
| **Moss** | 6 |  | 6 | 10 |  | 10 | 10 | 2 | 12 | 28 |
| *Distichium capillaceum* | 1 |  | 1 | 6 |  | 6 | 2 | 1 | 3 | 10 |
| *Stegonia latifolia* | 1 |  | 1 |  |  |  | 2 |  | 2 | 3 |
| *Tortella alpicola* | 1 |  | 1 | 1 |  | 1 |  |  |  | 2 |
| *Bryum argenteum* |  |  |  |  |  |  |  | 1 | 1 | 1 |
| *Bryum* spp. | 2 |  | 1 | 1 |  | 1 | 4 |  | 4 | 5 |
| *Didymodon* spp. |  |  |  |  |  |  | 1 |  | 1 | 1 |
| *Myurella julacea* |  |  |  | 1 |  | 1 |  |  |  | 1 |
| *Orthothecium strictum* |  |  |  | 1 |  | 1 |  |  |  | 1 |
| *Buckia vaucheri* |  |  |  |  |  |  | 1 |  | 1 | 1 |
| *Syntrichia* spp. | 1 |  | 1 |  |  |  |  |  |  | 1 |
| **Plant** | 15 | 8 | 23 | 27 | 8 | 35 | 26 | 10 | 36 | 94 |
| *Carex borii* |  |  |  | 8 |  | 8 | 3 |  | 3 | 11 |
| *Waldheimia tridactylites* | 3 | 2 | 5 | 1 |  | 1 |  | 3 | 3 | 9 |
| *Thylacospermum caespitosum* |  | 3 | 3 | 2 | 2 | 4 |  | 1 | 1 | 8 |
| *Saussurea glacialis* |  | 1 | 1 | 3 | 2 | 5 |  | 1 | 1 | 7 |
| *Carex sagaensis* |  |  |  | 2 |  | 2 | 4 |  | 4 | 6 |
| *Potentilla pamirica* |  |  |  |  | 1 | 1 | 5 |  | 5 | 6 |
| *Sibbaldia tetrandra* |  |  |  | 3 |  | 3 | 2 |  | 2 | 5 |
| *Poa attenuata* |  | 1 | **1** | 1 |  | 1 |  | 3 | **3** | 5 |
| *Carex nivalis* | 4 |  | **4** |  |  |  |  |  |  | 4 |
| *Kobresia schoenoides* |  |  |  | 1 |  | **1** | 3 |  | **3** | 4 |
| *Leontopodium ochroleucum* |  |  |  | 1 |  | **1** | 3 |  | **3** | 4 |
| *Waldheimia nivalis* | 3 |  | **3** |  |  |  |  |  |  | 3 |
| *Astragalus confertus* |  |  |  | 1 |  | **1** | 2 |  | **2** | 3 |
| *Arenaria bryoides* |  |  |  |  | 2 | **2** |  |  |  | 2 |
| *Bistorta vivipara* |  |  |  |  |  |  | 1 |  | **1** | 1 |
| *Dasiphora dryadanthoides* |  |  |  | 1 |  | **1** |  |  |  | 1 |
| *Draba altaica* | 1 |  | **1** |  |  |  |  |  |  | 1 |
| *Draba lasiophylla* | 1 |  | **1** |  |  |  |  |  |  | 1 |
| *Elymus jacquemontii* |  |  |  |  |  |  | 1 |  | **1** | 1 |
| *Kobresia pygmaea* |  |  |  |  | 1 | **1** |  |  |  | 1 |
| *Lloydia serotina* |  |  |  | 1 |  | **1** |  |  |  | 1 |
| *Nepeta longibracteata* | 1 |  | **1** |  |  |  |  |  |  | 1 |
| *Oxytropis platysema* |  |  |  |  |  |  | 1 |  | **1** | 1 |
| *Saussurea gnaphaloides* |  |  |  |  |  |  |  | 1 | **1** | 1 |
| *Saxifraga nanella* |  |  |  |  |  |  |  | 1 | **1** | 1 |
| *Saxifraga pulvinaria* |  |  |  | 1 |  | **1** |  |  |  | 1 |
| *Silene gonosperma* | 1 |  | **1** |  |  |  |  |  |  | 1 |
| *Stellaria decumbens* |  | 1 | **1** |  |  |  |  |  |  | 1 |
| *Tanacetum spp.* |  |  |  | 1 |  | **1** |  |  |  | 1 |
| *Thalictrum alpinum* |  |  |  |  |  |  | 1 |  | **1** | 1 |
| *Urtica hyperborea* | 1 |  | **1** |  |  |  |  |  |  | 1 |

**Supplementary Table 2.**
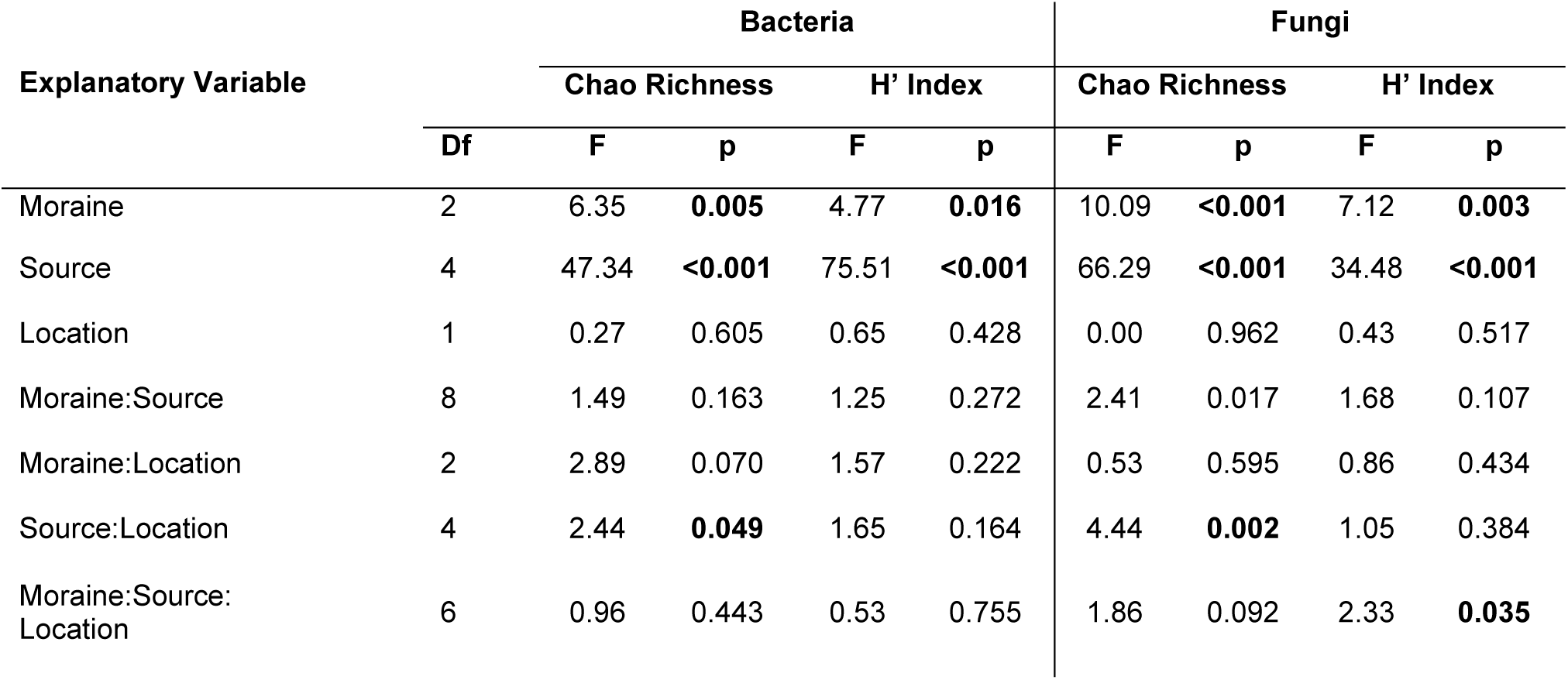
Analysis of variance (Type II F-test) results for alpha diversity metrics of fungal and bacterial communities across categorical moraine ages and different sources. Models were fit using linear mixed-effects models with Plot ID included as a random effect. Significance was assessed using Type II F-tests to evaluate the contribution of fixed effects with corrected degrees of freedom.

**Supplementary Table 3.**
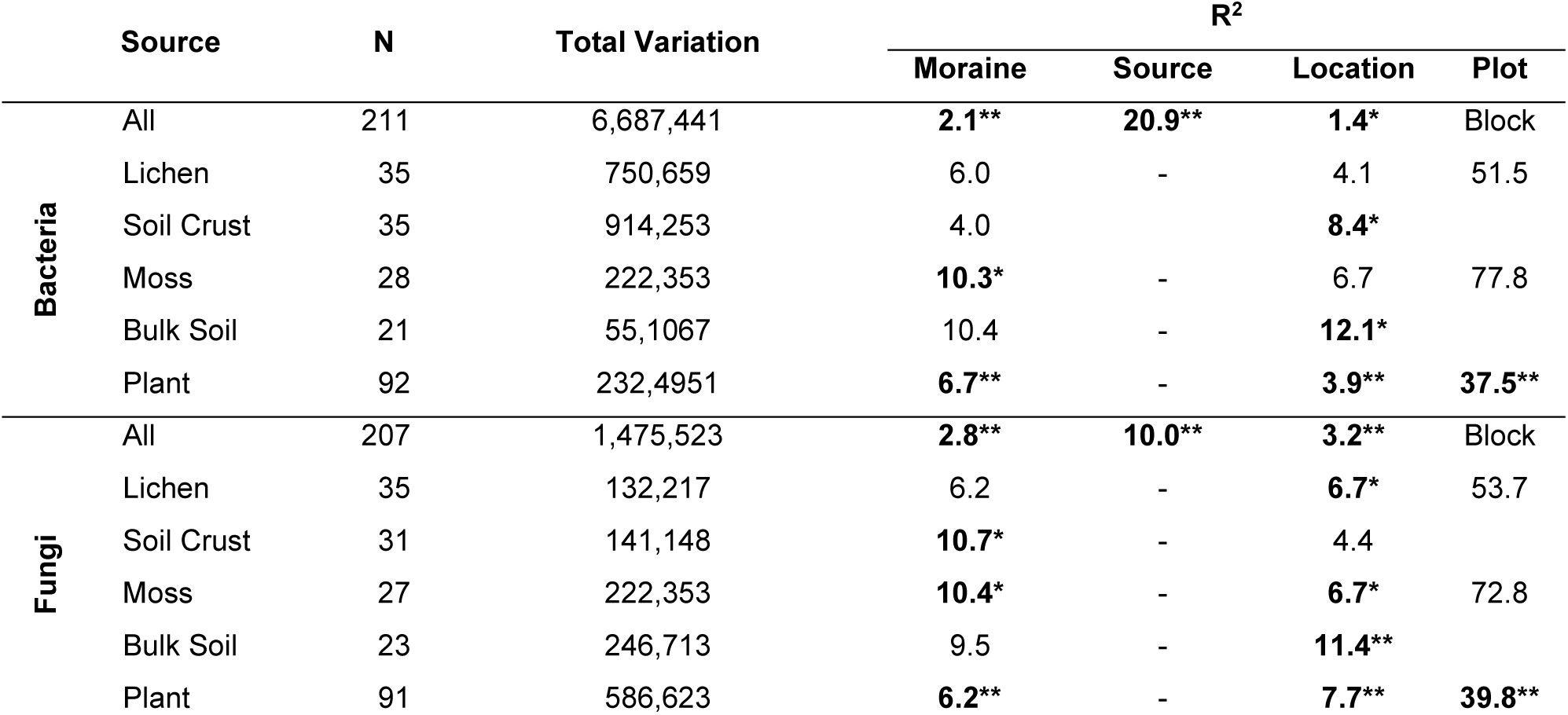
PERMANOVA results for bacterial and fungal community composition across moraines of different categorical ages and locations (Nubra and Tso Moriri), analyzed for all samples combined and separately by source. Community data were CLR-transformed and Euclidean distances were used for the analysis. R² values indicate the proportion of variance explained. Significance is indicated by asterisks (*p_adj_ < 0.05, ** p_adj_ < 0.01). Plot was included as a blocking factor in the combined analysis, and as a fixed factor in separate analyses when applicable.

**Supplementary Table 4.**
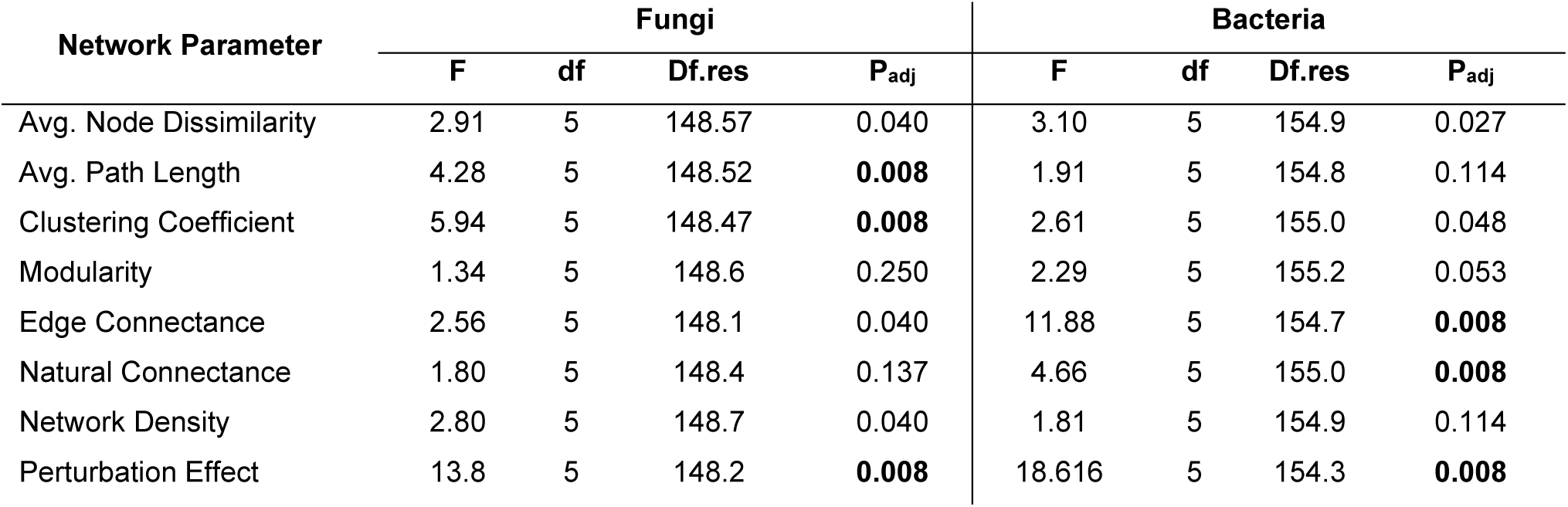
ANOVA results for global network parameters derived from bacterial and fungal community networks, assessed across separate source exclusions. Data were analyzed non-parametrically using an aligned rank transformation approach, followed by an ANOVA with plot ID as a random factor. with p-values adjusted using false discovery rate (FDR) correction. Bolded values indicate FDR-corrected p-values < 0.01, which are visualized in boxplots in Figure 4.

**Supplementary Figure 1.**
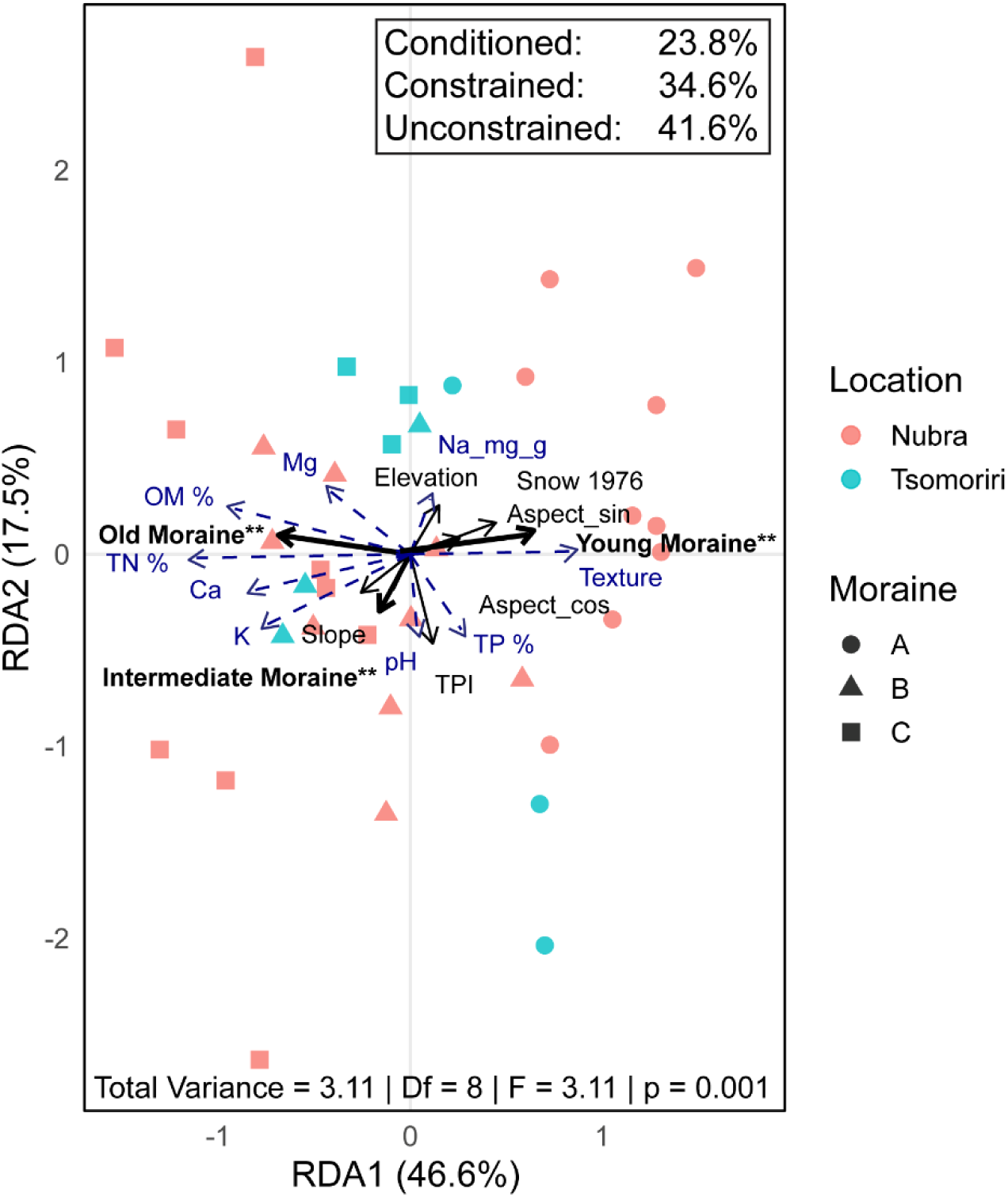
Redundancy analysis (RDA) of soil physicochemical measurements as response variables and environmental parameters as explanatory variables across four glacier forefields in Ladakh, India. Soil variables were RMS-scaled prior to analysis. The model was conditioned by “Location” to account for spatial structure among sites.

**Supplementary figure 2.**
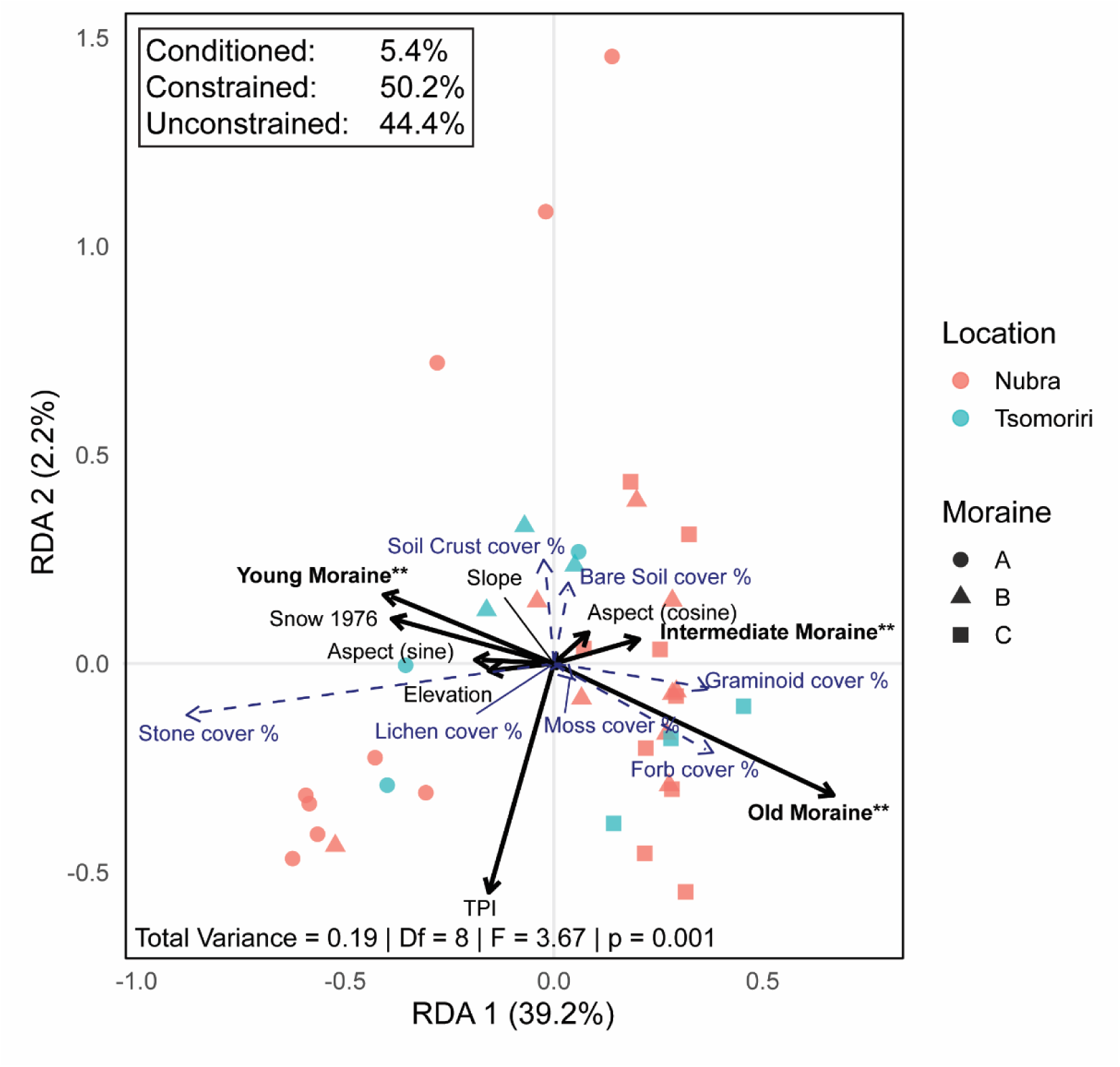
Redundancy analysis (RDA) of plot cover measurements—including plants, moss, lichen, soil crust, bare soil, and stones—as response variables, with environmental parameters as explanatory variables, across four glacier forefields in Ladakh, India. Cover data were transformed using the isometric log-ratio (ILR) transformation. The model was conditioned by “Location” to account for spatial structure among sites.

